# Combined High Fat and Viral Insults in Pregnancy Disrupt Specific Placental Nutrient Transporter Systems in Mice

**DOI:** 10.1101/2025.10.28.685153

**Authors:** Thaina Ferraz, Lucas Cardoso, Sadra Mohammadkhani, Enrrico Bloise, Kristin L Connor

**Author notes:** Correspondence: Dr. Kristin Connor, Department of Health Sciences, Carleton University, Ottawa, Ontario, Canada, K1S5B6.

## Abstract

The placenta adapts to support nutrient transport to meet the demands of the growing fetus. Maternal obesity and viral infection are associated with adverse fetoplacental development. How co-exposure affects placental morphology and nutrient transport across gestation remains unclear. We hypothesise that co-exposure would have a greater impact on placental morphology and nutrient transporter expression than either of the exposures alone. C57BL/6J mice were fed a control (CON) or high-fat (HF) diet before and throughout pregnancy and exposed to poly(I:C) or vehicle at gestational days (GD) 12.5, 15.5, or 18.5. Placental morphology, folate and fatty acid transporter mRNA expression and localisation were evaluated by haematoxylin and eosin staining, immunohistochemistry (reported as % immunoreactivity (ir)-area stained), and qPCR, respectively. At GD12.5, HF diet associated with reduced junctional zone (JZ) area and increased maternal blood space, followed by reduced fetal blood space at GD15.5 and increased JZ and labyrinth zone (LZ) area size by GD18.5. HF diet and viral infection independently associated with increased placental *Fat/Cd36* mRNA expression at GD12.5. However, HF diet and infection associated with increased ir-FR-α while HF diet alone decreased ir-PCFT at GD12.5. By GD15.5, HF diet associated with reduced placental ir-FR-α but increased ir-PCFT. At GD18.5, infection associated with reduced *Rfc1* mRNA expression. HF diet and infection associated with greater prevalence of SGA and LGA fetuses, particularly at GD18.5. Obesity and infection independently and jointly disrupted placental morphology and nutrient transporter expression in a zone- and gestational age-specific manner, with consequences for fetal growth across gestation.

## Introduction

Lifelong health is shaped by growth and development in early life, of which the placenta plays an essential role. Proper placental development and function are essential for meeting both the metabolic demands of the placenta itself, and those of the developing fetus^2^. Further, the placenta must continually adapt its structure and transport capacity to meet increasing fetal nutritional requirements throughout gestation^1^. However, these adaptations can be compromised by adverse maternal conditions, such as malnutrition^2^ and infection^3^, which may impair placental nutrient transfer and thus, affect fetal development^4^. Although maternal malnutrition^5^ and viral infection^6^ are highly prevalent worldwide, and are each associated with adverse pregnancy outcomes including placental dysfunction and impaired fetal growth^7-10^, there is limited data across pregnancy on how the placenta develops, maintains, and regulates nutrient transport in response to these conditions, especially when they co-occur.

Some of the most important nutrients for fetoplacental development include fatty acids and folate^11,12^. Fatty acids support placental angiogenesis in early development, and provide energy for placental growth, ensuring the development of key placental structures such as the chorionic villi^13^. Folate, a central component of one-carbon metabolism, supports the synthesis of purines and pyrimidines and the methylation of DNA, RNA, and proteins^14^, and supports cell proliferation^15^. These factors are necessary for placental tissue growth^16^, especially in early pregnancy when folate contributes to the development of chorionic villi and vascular remodelling required for efficient nutrient and gas exchange at the maternal-fetal interface^17,18^. Placental transporters responsible for the uptake and transfer of fatty acids and folate are differentially expressed throughout gestation and across placental regions, reflecting their essential roles in supporting placental function and fetal development^11,12^. In mice, placental fatty acid transporters are expressed during mid-to-late gestation (approximately GD11.5-18.5), with fatty acid transporters predominantly localised to the placental labyrinth zone (LZ; exchange area between mother and fetus) and decidua^12^. Folate transporters are localised in glycogen trophoblast cells within the placental junctional zone (JZ; the endocrine region)^11^ with the earliest expression of folate transporters in mouse placenta reported at GD8.5, and present until term at GD18.5^11^.

Beyond their role in placental development, the efficient transplacental transfer of key nutrients such as fatty acids and folate is essential for optimal fetal growth and development ^19-21^. Fatty acids participate in cellular signalling and are indispensable for fetal organogenesis^13,22^, placental and fetal lipid metabolism, required for growth and development^13^, fetal vascular formation and brain myelination throughout gestation^23^. In humans, maternal lipid levels measured at 13 weeks of gestation have been shown to modulate fetal growth rates, potentially exerting long-term effects on offspring metabolic health^24,25^. Additionally, excessive fetal exposure to fatty acids has been shown to contribute to abnormal growth trajectories, such as increased placental weight and increased fetal fat deposition^26^, shown by elevated levels of cholesterol and low-density lipoprotein in newborns^27^Folate is crucial for embryonic haematopoiesis and neural tube closure^16^. Environmental or nutritional factors that diminish maternal or fetal folate availability may disrupt one-carbon metabolism, impair fetal growth and increase the risk neural tube defects and altered fetal development^14^. Notably, reduced maternal folate levels in early human pregnancy have been associated with smaller placental volume, which can compromise placental function and thereby restrict fetal development^28^. Together, fatty acids and folate play pivotal roles in shaping fetal growth trajectories and developmental outcomes^1^.

Since effective maternal-to-fetal transport of these nutrients is mediated by specific placental transporters, any disruption to transporter expression or function could have downstream effects on fetoplacental development. We have shown that maternal high fat (HF) diet exposure, permissive of obesity and viral infection in mice can independently impair placental development and function^29^, and may do this through disrupting placental morphology and selective nutrient exchange^30^. In humans, obesity has been shown to increase both systemic and placental expression of pro-inflammatory markers including elevated levels of maternal plasma monocyte chemoattractant protein-1 (MCP-1) and tumour necrosis factor-alpha (TNF-α), with increased placental expression of interleukin-1 beta (IL-1β), and activation of the p38 mitogen-activated protein kinase (p38-MAPK) and signal transducer and activator of transcription 3 (STAT3) inflammatory pathways^31^. Additionally, placental fatty acid transporters, such as fatty acid transporter protein 2 (FATP2) and 4 (FATP4)^26^, can enhance the transfer of both low-grade inflammatory signals and lipids to the fetus. In contrast, during first and second trimester of human pregnancies, maternal obesity has been associated with a reduction in protein expression of placental folate transporters, including folate receptor α (FR-α) and reduced folate carrier 1 (RFC1)^32^, potentially limiting fetal folate availability and compromising fetal development and growth^32^. Moreover, placental folate receptor beta (*Fr-β*) expression has been shown to be increased by late gestation in pregnancies were dams had obesity compared undernourished dams^33^. These findings suggest that obesity creates a transfer imbalance in key nutrients that could compromise fetoplacental development and growth at different gestational ages. There is much less known about the effects of viral infection on placental nutrient transport. In the context of viral infection, infection-induced lipid droplet production, which supports viral survival, depends on FATP and fatty acid binding protein (FABP)^34^ expression. Additionally, a study examining placentae from women with chronic hepatitis C, showed increased placental RFC mRNA expression^35^. Yet, no studies have specifically investigated how acute viral infection alone, or associated with maternal obesity, alters placental fatty acid and folate transporter expression.

Using an animal model, we hypothesised that maternal HF diet permissive of obesity and acute viral infection would each independently alter placental morphometry and fatty acid and folate transporter expression, resulting in altered fetoplacental growth, in a gestational-age dependent manner. We further hypothesised that co-exposure to high fat diet and viral infection would exacerbate the isolated effects of both stressors, leading to greater disruption of placental morphology, transporter function, and fetoplacental growth. Understanding how placental morphometry, fatty acid and folate transporter systems are regulated at the maternal-fetal interface across pregnancy, may provide critical insights into the mechanisms by which maternal obesity and viral infection (individually or in combination) compromise pregnancy outcomes.

## Methods

### Animal model and study design

All animal procedures were conducted in accordance with a protocol approved by the Carleton University Animal Care Committee (AUP 114994). Further, methodology and results were reported in accordance with the ARRIVE guidelines (Supplementary Table 1)^36^. The experimental model has been reported previously^29^. Male and female C57BL/6J mice, aged five to six weeks, were maintained in the same housing room at 25°C on a 12h light/12h dark schedule, with unrestricted access to food and water. Prior to breeding, females were randomly allocated to either a high fat (HF; n=28) or control (CON; n=34) dietary intervention. The HF group received Teklad custom diet, permissive of obesity, TD.210535 (62% fat, 4% inulin, 18.8% protein, 19.2% carbohydrates; energy density: 5 kcal/g), whereas the CON group received Teklad custom diet TD.210534 (15.8% fat, 4% inulin, 20.9% protein, 63.3% carbohydrates; energy density: 3.5 kcal/g; Inotiv, Indianapolis, IN, USA). Diets were provided *ad libitum* beginning six to eight weeks before mating and were maintained throughout gestation. At each of three gestational ages (GD12.5, GD15.5, and GD18.5), dams within each dietary groups were further assigned to receive a single intraperitoneal injection of either the double-stranded RNA, Poly(I:C) (10 mg/kg), which is known as an acute viral-like immune challenge or saline vehicle (0.9%; 0.1 mL). Injections were given 24 h before tissue collection. Further, each gestational time point included four groups: CON-VEH, CON-POLY, HF-VEH, and HF-POLY, with 5-9 dams per group. Gestational time points were chosen to assess placental development and function at three key developmental stages: GD12.5, when the mouse placenta has developed its major structural compartments^37^; GD15.5, when the placenta reaches functional maturity^37^ and main placental structures undergo further development and structural changes to optimise nutrient exchange through the end of pregnancy^29^. Additionally, between GD15.5 and 18.5 the placenta mobilises glycogen stores to support fetal energy demands and prepare for delivery^37,38^. The Poly(I:C) dose of 10 mg/kg was determined based on a pilot study in pregnant C57BL/6J mice, as previously described^29^.

### Biospecimen collection and processing

Dams were euthanised via cervical dislocation at GD12.5, 15.5 and 18.5, after which trunk blood was collected into heparinized tubes. The uterus was immediately dissected to isolate fetuses and placentae, which were weighed and processed either by flash-freezing in liquid nitrogen for molecular analyses or fixed in 10% neutral-buffered formalin for 48 hours for histological evaluation. Fetal tails obtained from tissue collection were used to determine the sex of each fetus by PCR genotyping. DNA was extracted from the tail samples with the Extract-N-Amp Tissue PCR Kit (Sigma-Aldrich, Oakville, ON, Canada) according to the supplier’s instructions. Sex-specific chromosome sequences were then amplified by PCR, and the resulting products were identified following separation on an agarose gel electrophoresis. Primer sequences are provided in Supplementary Table 2. One male and one female placenta per litter were selected from each dietary and infection group, based on mean placental weight per group, for subsequent assays. Fetal and placental weight distributions were analysed across gestation and by experimental group. At each GD, fetuses were classified as small-(SGA; <10th percentile), appropriate-(AGA; 10^th^-90th percentile), or large-for-gestational-age (LGA; >90th percentile) based on percentile-based cutoffs according to the overall weight distribution by GD.

### Histological analyses

Paraffin-embedded placental samples were sectioned at 5 µm from the midline of the placenta and stained with haematoxylin solution, Gill’s No. 1 and eosin Y (H&E; Thermo Fisher; Ottawa, ON, Canada) according to standard protocols. Placental H&E images were captured at 4x magnification using the Invitrogen EVOS FL Auto 2 Imaging System (version: 2.0.2094.0, Thermo Fisher Scientific; Ottawa, ON, Canada). In each field of view, we captured full area of placental junctional and labyrinth zones (LZ; the nutrient exchange region). The boundaries of the LZ and JZ zones were manually delineated in ImageJ (version 1.54d) by a single observer, blinded to experimental group, in three different slides and averaged per placentae. Quantification of absolute LZ and JZ areas (µm^2^) was obtained by multiplying pixel counts by the calibrated pixel area, and relative areas were expressed as the fraction of total placental sectional area. Scale calibration was performed using a stage micrometer (1µm/pixel). Further, in a subset of placentae, maternal and fetal blood space areas were quantified in ImageJ using a grid overlay for point counting. For each placenta, three non-overlapping fields of view were captured at 40x magnification, and the count of points falling within maternal or fetal blood spaces was recorded and averaged per placentae. Interhaemal membrane thickness was measured at 40x magnification using an orthogonal intercept length in 30 random sections planes in three non-overlapping placental fields of view and averaged per placentae.

### Immunohistochemistry

Next, we evaluated whether and how maternal HF diet and viral infection exposure affected fatty acid and folate transporter expression, we performed IHC as previously described^29^. In brief, serial 5 µm paraffin-embedded placental sections were obtained from the midline region of one male and one female placenta per litter and stained to localise and semi-quantitatively assess immunoreactive (ir)-area stained for FR-α, PCFT (protein coupled folate transporter) and RFC1. Primary antibody incubation for FR-α (1:250; ab67422, Abcam; Boston, MA, USA), PCFT (1:500; NBP1-06603, Novus Biologicals; Toronto, ON, Canada) and RFC1 (1:200; ab62302, Abcam; Boston, MA, USA), occurred at 4°C overnight. Next, biotinylated goat anti-rabbit (Vector BA-1000; Brockville, ON, Canada) in antibody diluent (DAKO; Mississauga, ON, Canada) was applied for 1 hour at room temperature (RT). For each placenta, three randomly selected, non-overlapping fields of view were imaged at 20x magnification using the Invitrogen EVOS FL Auto 2 Imaging System (version: 2.0.2094.0, Thermo Fisher Scientific; Ottawa, ON, Canada). Images were acquired within the placental LZ and JZ under identical exposure conditions. Target protein localisation (cell type and spatial distribution within LZ and JZ) was evaluated qualitatively, and the ir-stained area was quantified semi-quantitatively using ImageJ software (version: 1.54d) for FR-α, PCFT and RFC1. As previously described^29^, for each field of view, ir was assessed by determining the percentage of tissue exhibiting positive staining. For each placenta, ir measurements were obtained from three separate fields and averaged to generate a mean percentage (%) of ir-area stained. All protein staining assessments were performed by the same investigator, who remained blinded of group allocation during the analysis.

### RNA isolation and expression in the placenta

Placental total RNA was isolated using the RNeasy Mini Kit (QIAGEN), following previously reported procedures^2,29^. RNA quantity and purity were determined by NanoDrop spectrophotometry, while RNA integrity was confirmed using gel electrophoresis. For cDNA synthesis, 1 µg of RNA meeting the required quality criteria was reverse transcribed using 5X iScript Reverse Transcription Supermix (Bio-Rad, Mississauga, ON, Canada). Samples prepared without reverse transcriptase (NRT) served as negative controls in subsequent PCR assays. Gene expression was subsequently quantified by real-time PCR using a Bio-Rad CFX384 system (Hercules, CA, USA). The fatty acid-related genes examined included: *Fatp1, Fatp4* (fatty acid transport proteins 1 and 4), *Fat/Cd36* (fatty acid translocase) *Fabp*_*pm*_ (fatty acid binding protein plasma membrane) and folate genes: *Fr-α, Pcft, Rfc1* and *Fr-β*. Primer sequences for genes of interest and three stably expressed reference genes (*β-actin, Ywhaz, Gapdh*) are listed in Supplementary Table 2. SsoAdvanced Universal SYBR Green Supermix (Bio-Rad, Mississauga, ON, Canada) was used with 0.8 µM concentration of gene-specific forward and reverse primers. Standard curves were established for each target gene to confirm amplification efficiency. All standards, samples, and negative controls were analysed in triplicate following an initial denaturation at 95°C for 20s, 40 cycles of 95°C for 5s and 60°C for 20s. Melt curve analysis occurred from 65°C to 95°C, with 0.5°C increments every 5s. Relative transcript concentration was determined using the comparative ΔΔCq method, where target gene Cq values were normalised to the geometric mean of reference genes to obtain ΔCq, and fold-change expression was calculated relative to the control group (ΔΔCq)^39^

### Statistics

All data were assessed for normality (Shapiro–Wilk test) and when required, non-normal values were log-transformed to achieve normality. Homogeneity of variances was evaluated (Levene’s test). When a linear mixed model (LMM) was used to evaluate study outcomes (described below), heterogeneity of variance was incorporated into the model structure, including potential litter effects. To evaluate the main fixed effects of diet and infection a LMM analysis was performed for each GD (12.5, 15.5, and 18.5), between groups (CON-VEH, CON-POLY, HF-VEH, HF-POLY) for maternal and placental measures with litter included as a random effect. Placental and fetal outcomes were also stratified by fetal sex and analysed by Two-way Anova. Post hoc comparisons between groups were assed using Tukey’s test for all parametric analyses. Data that could not be normalized by log transformation were assessed using the Kruskal-Wallis test, followed by Steel-Dwass post hoc. Normally distributed data are reported as mean ± standard deviation, whereas non-normally distributed data is summarized using the median and interquartile range (IQR). Figures display diamonds representing 95% confidence intervals (CIs). When transformations were required for statistical testing, results are shown on their original, untransformed scale. A p-value ≤0.05 was considered statistically significant.

Standardized effect size measures such as Cohen’s d are not appropriate for linear mixed models (LMMs) or non-parametric analyses in which Hodges-Lehmann (HL) estimates are reported^40^. Therefore, for LMM analyses, fixed-effect estimates (B coefficients) are presented alongside their standard errors and 95% confidence intervals (CIs). For maternal and placental outcomes that did not meet normality assumptions and could not be normalized through log transformation, HL estimates of the median difference between groups with corresponding 95% CIs are reported. JMP statistical software (Version:18.0.2) was used to for data analyses.

## Results

### Maternal HF diet impacts placental architecture throughout gestation

Overall, differences in placental architecture were influenced mostly by maternal HF diet. First, at GD12.5, HF diet was associated with decreased JZ (Figure 1A), but not LZ (Figure 1D), area compared to CON (HL=−528.865 95% CI [−928.99, −223.75], p=0.0008). At GD15.5, neither diet, infection nor co-exposure of maternal diet and infection influenced JZ or LZ area. At GD18.5, maternal HF diet was associated with increased JZ (B=−0.05 [−0.103, −0.013], p=0.01; Figure 1C) and LZ (B=−0.04 [−0.08, −0.004], p=0.03; Figure 1F) areas compared to CON-fed dams. Neither infection nor co-exposure of maternal diet and infection influenced JZ or LZ area at any GD.

**Figure 1.**
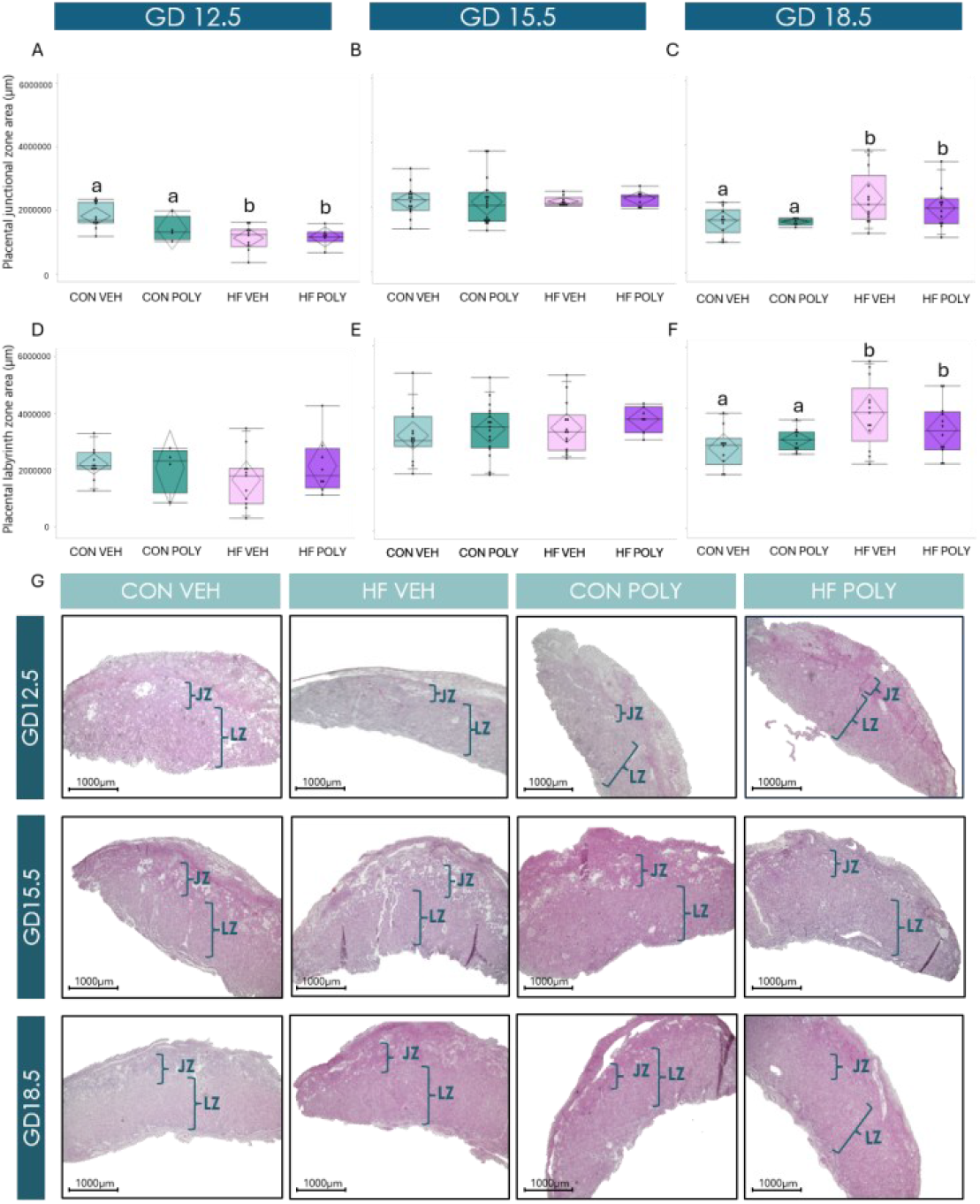
Effect of maternal diet and infectious exposure on placental JZ and LZ area across gestation. **A**. Placental JZ area stratified by dietary and infectious exposures at GD12.5. Maternal HF diet was associated with decreased placental JZ area size compared to CON (p=0.0008). **B**. Placental JZ area stratified by dietary and infectious exposures at GD15.5. **C**. Placental JZ area stratified by dietary and infectious exposures at GD18.5. Maternal HF diet was associated with increased placental JZ area size compared to CON (p=0.006). **D**. Placental LZ area stratified by dietary and infectious exposures at GD12.5. **E**. Placental LZ area stratified by dietary and infectious exposures at GD15.5. **F**. Placental LZ area stratified by dietary and infectious exposures at GD18.5. Maternal HF diet was associated with increased placental LZ area size compared to CON (p=0.008). Data are quantile box plots with 95% CI diamonds. Linear mixed model and Tukey’s post hoc. Groups not connected by the same letter are significantly different, p≤0.05. **G**. Representative images of haematoxylin and eosin staining in mouse placentae showing JZ and LZ area. Scale = 1000μm, magnification = 4x. LZ = labyrinth zone JZ = junctional zone. GD = gestational day. CON = control. HF = high fat. VEH = vehicle. POLY = Poly(I:C). JZ = junctional zone. LZ = labyrinth zone. CI = confidence interval.

To explore whether the observed architectural alterations also influenced placental vascular compartments, which could impact nutrient transfer to the fetus, we next assessed placental interhaemal space and maternal and fetal blood space area in a subset of placentae. At GD12.5 neither HF diet, infectious exposure, nor their co-exposure impacted placental interhaemal space thickness or fetal blood space area (Figures 2A, G). However, at GD12.5 maternal HF diet, but not infection, was associated with increased maternal blood space area compared to CON (B = −2.34 [−4.70, 0.01], p=0.05; Figure 2D), which did not persist at later GDs (Figure 2E, F). Importantly, at GD15.5, maternal HF diet alone was associated with reduced fetal blood space area compared to CON (B = 0.09, [0.001, 0.182], p=0.04; Figure 2H), with no further changes induced by maternal HF diet and/or infectious exposure observed on placental interhaemal space thickness or maternal blood space area (Figures 2B, E). Additionally, neither HF diet, infectious exposure, nor their co-exposure impacted placental interhaemal space or maternal and fetal blood space area at GD18.5 (Figures 2C, F, I).

**Figure 2.**
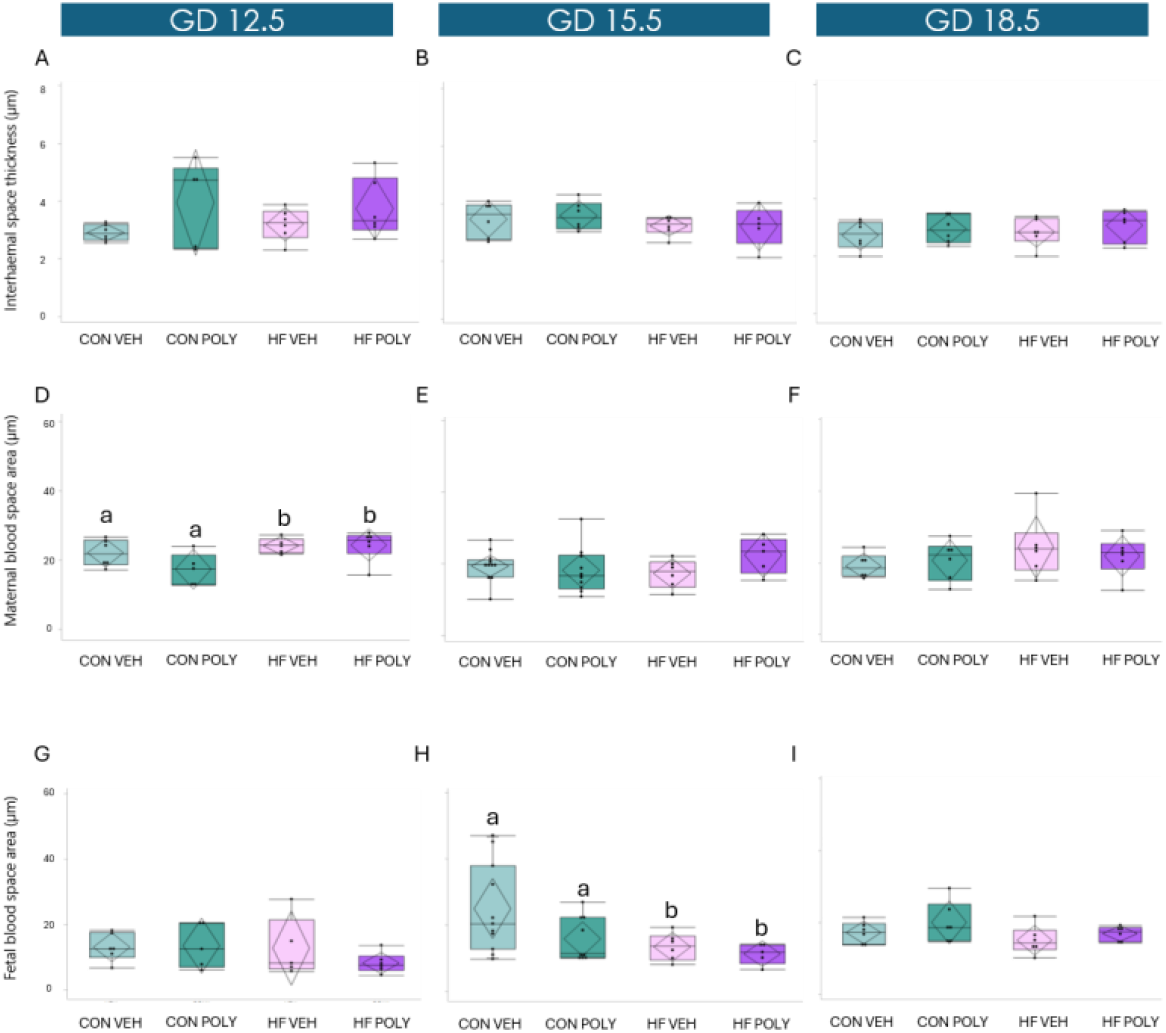
Effect of maternal diet and infectious exposure on placental interhaemal and blood space area across gestation. **A**. Placental interhaemal space stratified by dietary and infectious exposures across gestation. **A, B, C**. Neither HF diet and/or Poly(I:C) was associated with altered interhaemal space in any GD. **D**. Maternal blood space area stratified by dietary and infectious exposures at GD12.5. Maternal HF diet was associated with increased maternal blood space area compared to CON (p=0.05). **E, F**. Neither HF diet and/or Poly(I:C) was associated with altered maternal blood space area at GD15.5 (E) or GD18.5 (F). **H**. Fetal blood space area stratified by dietary and infectious exposures at GD15.5. Maternal HF diet was associated with reduced fetal blood space area compared to CON (p=0.04). **G, I**. Neither HF diet and/or Poly(I:C) was associated with altered fetal blood space area at GD12.5 (G) or GD18.5 (I). Data are quantile box plots with 95% CI diamonds. Linear mixed model and Tukey’s post hoc. Groups not connected by the same letter are significantly different p≤0.05. GD = gestational day. CON = control. HF = high fat. VEH = vehicle. POLY = Poly(I:C). CI = confidence interval.

### Maternal HF diet and infection independently impact placental Fat/Cd36 mRNA expression, but not other fatty acid transporters, at GD12.5

At GD12.5, HF diet was associated with increased placental mRNA expression of *Fat/Cd36* compared to CON (B=−0.16 [−0.29, −0.03], p=0.01; Figure 3G), while acute infection alone was also associated with increased *Fat/Cd36* mRNA expression levels compared to VEH (B=−0.16 [−0.29, −0.02], p=0.02; Figure 3G). Co-exposure to HF diet and infection did not further impact *Fat/Cd36* mRNA expression levels at GD12.5 (Figure 3G). Overall, HF diet and/or viral infection were not associated with changes in placental mRNA expression levels of *Fatp1, Fatp4* or *Fabp*_pm_, at any GD (Figure 3).

**Figure 3.**
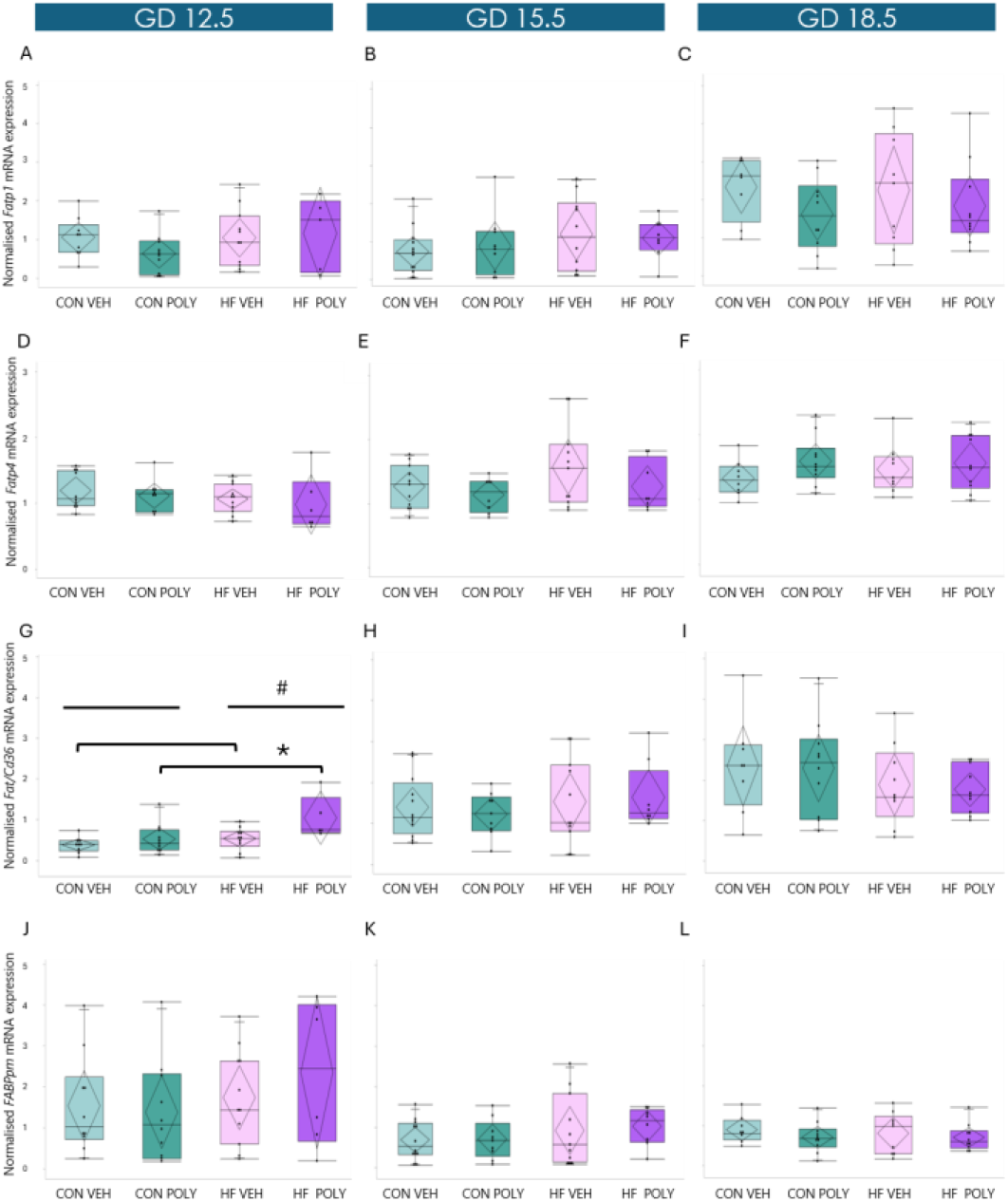
Effect of maternal diet and infectious exposure on placental fatty acid transporters mRNA expression across gestation. **A, B, C**. Placental *Fatp1* mRNA expression levels stratified by dietary and infectious exposures at GD12.5, 15.5 and 18.5. **D, E, F**. Placental *Fatp4* mRNA expression levels stratified by dietary and infectious exposures at GD12.5, 15.5 and 18.5. **G**. Placental *Fat/Cd36* mRNA expression levels stratified by dietary and infectious exposures at GD12.5. Maternal HF diet was associated with increased placental *Fat/Cd36* mRNA expression levels compared to CON (p=0.01). Poly(I:C) exposure was associated with increased placental *Fat/Cd36* mRNA expression levels compared to VEH (p=0.01). **H, I**. Placental *Fat/Cd36* mRNA expression levels stratified by dietary and infectious exposures at GD15.5 and 18.5. **J, K, L**. Placental *Fabp*_*pm*_ mRNA expression levels stratified by dietary and infectious exposures at GD12.5, 15.5 and 18.5. Data are quantile box plots with 95% CI diamonds. Linear mixed model and Tukey’s post hoc or Kruskall Wallis and Steel-Dwass post hoc. Groups not connected by the same letter are significantly different, #p≤0.05 vs. CON. *p≤0.05 vs. VEH. GD = gestational day. CON = control. HF = high fat. VEH = vehicle. POLY = Poly(I:C). CI = confidence interval.

### Maternal HF diet and infection differently impact placental folate transporter expression throughout gestation

Overall, HF diet and/or viral infection had no effect on placental mRNA expression levels of *Fr-α, Pcft* or *Fr-β* and at any GD (Figure 4). However, by GD18.5, maternal infection, but not HF diet, was associated with decreased placental mRNA expression levels of *Rfc1* compared to VEH (B =0.25 [0.02, 0.49], p=0.03; Figure 4I). Co-exposure to HF diet and infection did not further impact placental mRNA expression levels of *Rfc1* (Figure 4I).

**Figure 4.**
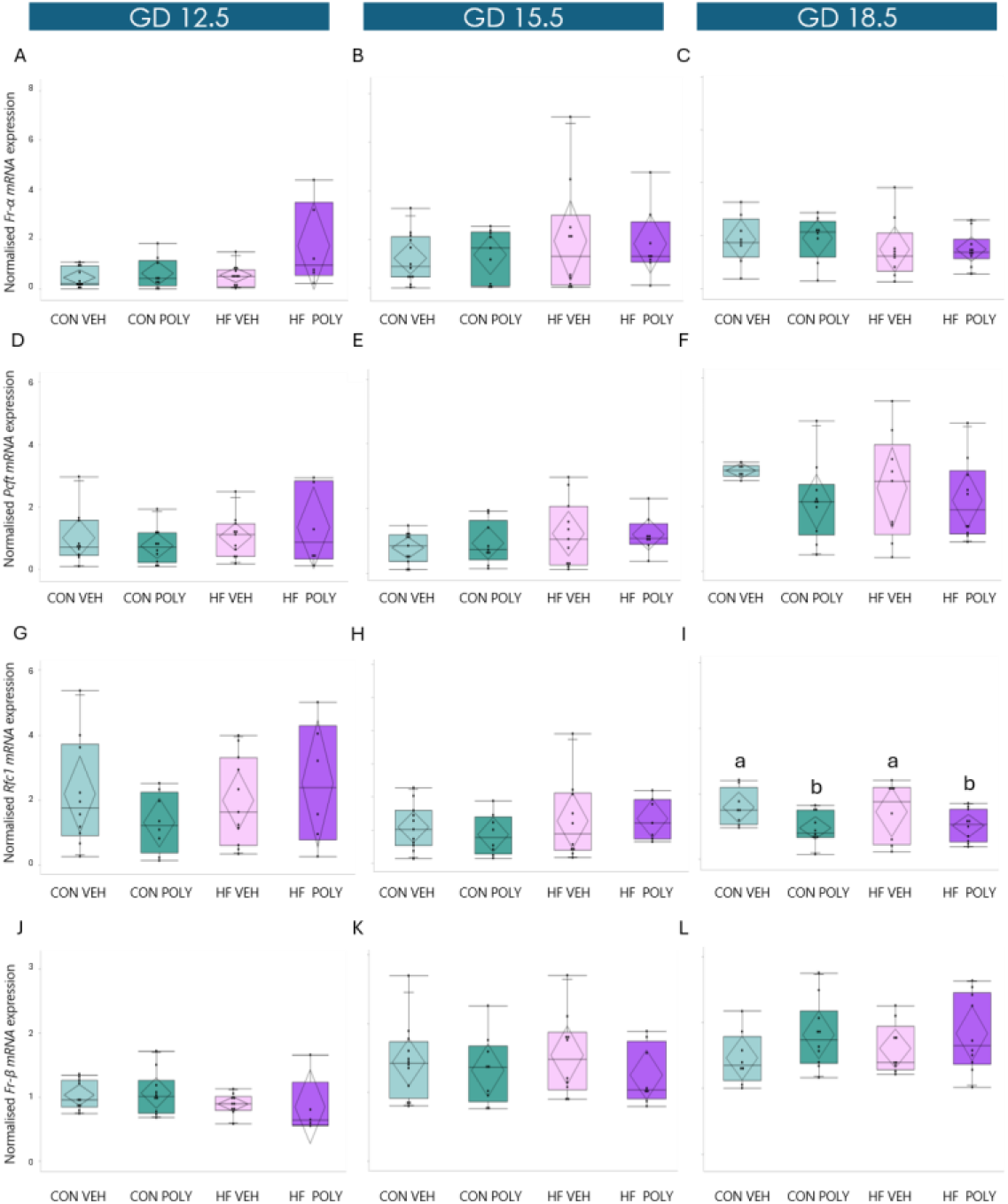
Effect of maternal diet and infectious exposure on placental folate transporters mRNA expression across gestation. **A, B, C**. Placental *Fr-α* mRNA expression levels stratified by dietary and infectious exposures at GD12.5, 15.5 and 18.5. **D, E, F**. *Pcft* mRNA expression stratified by dietary and infectious exposures at GD12.5, 15.5 and 18.5. **G, H**. Placental *Rfc1* mRNA expression levels stratified by dietary and infectious exposures at GD12.5 and 15.5. **I**. Placental *Rfc1* mRNA expression stratified by dietary and infectious exposures at GD18.5. Poly(I:C) exposure was associated with decreased placental *Rfc1* mRNA expression levels compared to VEH (p=0.03). **J, K, L**. Placental *Fr-β* mRNA expression levels stratified by dietary and infectious exposures at GD12.5, 15.5 and 18.5. Data are quantile box plots with 95% CI diamonds. Linear mixed model and Tukey’s post hoc. Groups not connected by the same letter are significantly different, p≤0.05. GD = gestational day. CON = control. HF = high fat. VEH = vehicle. POLY = Poly(I:C). CI = confidence interval.

We next evaluated localisation and % ir-area stained of placental folate transporters. Staining for FR-α, PCFT and RFC1 were both localised in the JZ and LZ across gestation, and they were mainly expressed in the JZ spongiotrophoblast (SpT) and LZ syncytiotrophoblast (STB) cytoplasm. At GD12.5, we observed that HF diet and infection associated with increased ir-FR-α % area stained in the JZ compared to VEH (B=4.62 [−0.06, 8.39], p=0.05; Figure 5A). At GD12.5 HF diet also associated with increased LZ ir-FR-α expression compared to CON (B =−21.50 [−41.17, −1.83], p=0.03; Figure 5D). Combined exposure of HF diet and infection had no impact on placental LZ ir-FR-α expression (Figure 5D). At GD15.5 in the JZ and LZ, maternal HF diet was associated with decreased ir-FR-α % area stained compared to CON (JZ: B=6.28 [0.41, 12.15], p=0.03; Figure 5B; LZ: B=7.26 [0.94, 13.58], p=0.02; Figure 5E). By GD18.5, maternal HF diet was associated with decreased ir-FR-α %area stained in the JZ compared to CON (B=9.39 [0.46, 18.32], p=0.04; Figure 5C), but not in the LZ (Figure 5F). Neither infection nor HF diet and infection further impacted ir-FR-α area stained in the JZ or LZ at GD15.5 or 18.5 (Figures 5B, C, E, F).

**Figure 5.**
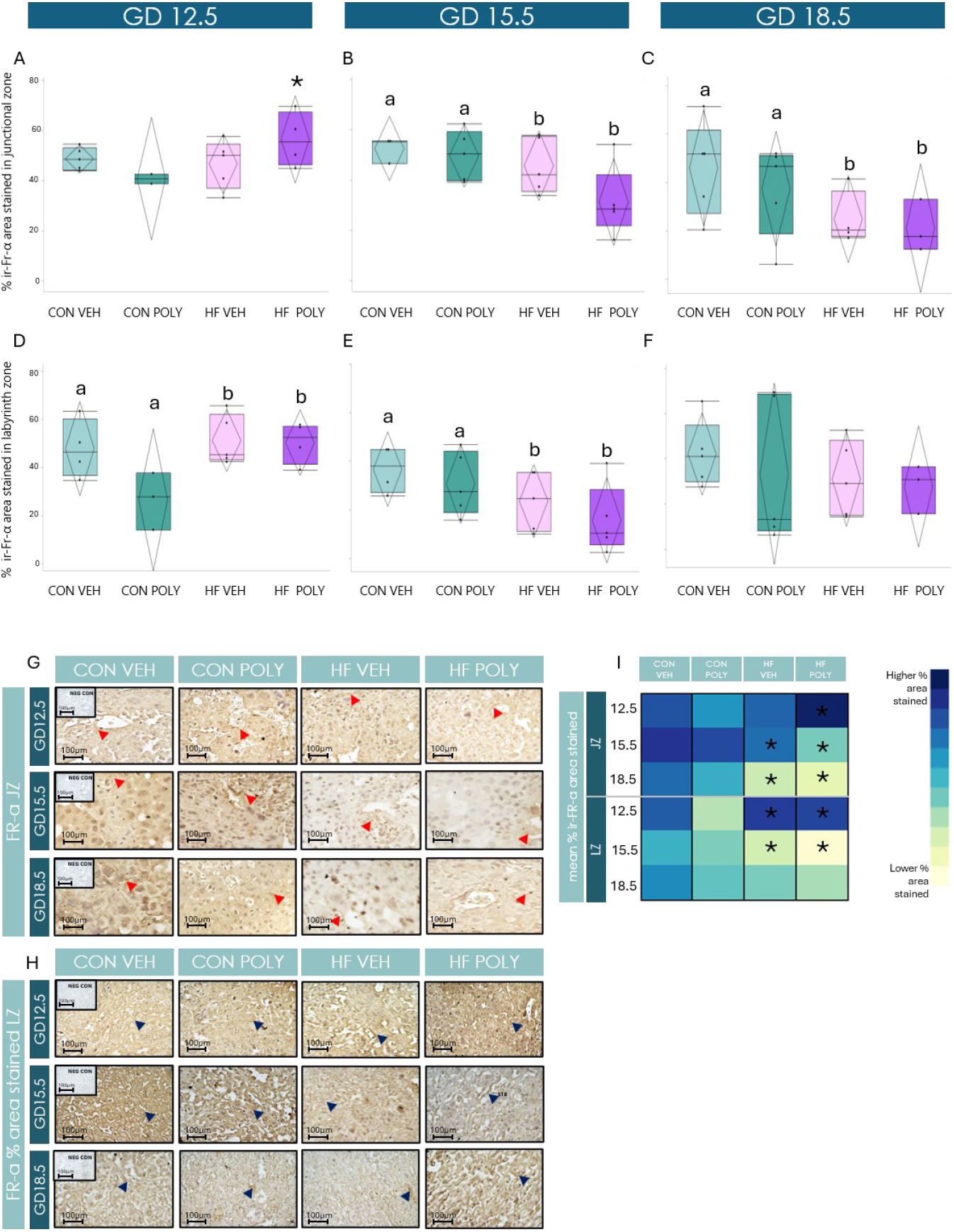
Effect of maternal diet and infectious exposure on placental ir-FR-α % area stained and localisation across gestation. **A**. Placental JZ ir-FR-α % ir-area stained stratified by dietary and infectious exposures at GD12.5. Maternal HF diet and infection was associated with increased ir-FR-α % ir-area stained in the JZ at GD12.5 compared to all other groups (p=0.05). **B, C**. Placental JZ ir-FR-α % ir-area stained stratified by dietary and infectious exposures at GD15.5 and 18.5. Maternal HF diet was associated with decreased ir-FR-α % ir-area stained in the JZ at GD15.5 (p=0.03) and 18.5 compared to CON (p=0.04). **D**. Placental LZ ir-FR-α % area stained stratified by dietary and infectious exposures at GD12.5. Maternal HF diet was associated with increased ir-FR-α % area stained in the LZ at GD12.5 compared to CON (p=0.02). **E**. Placental LZ ir-FR-α % area stained stratified by dietary and infectious exposures at GD15.5. Maternal HF diet was associated with decreased ir-FR-α % area stained in the LZ at GD15.5 compared to CON (p=0.03). **F**. Placental LZ ir-FR-α % area stained stratified by dietary and infectious exposures at GD18.5. Data are quantile box plots with 95% CI diamonds. Linear mixed model and Tukey’s post hoc. Groups not connected by the same letter are significantly different, p≤0.05. *Represents significant difference in HF POLY compared to all other groups. **G, H**. Representative images of ir-FR-α % ir-area stained in the JZ SpT (G) and LZ STB (H) cytoplasm across dietary and infectious exposures at different GD. Red arrows indicate positive FR-α immunostaining in JZ SpT. Blue arrows indicate positive FR-α immunostaining in LZ STB cytoplasm. Scale bar = 100μm, magnification = 40x. **I**. Heat map of mean % ir-area stained of Fr-α in the placental JZ and LZ by GD across exposure groups. Significant differences between dietary and infectious groups in different placental zones and at different GD (p≤0.05) are marked with an asterisk. Darker blue = higher % ir-area stained; Lighter blue/yellow = lower % ir-area stained. CON. GD= gestational day. CON = control. HF = high fat. VEH = vehicle. POLY = Poly(I:C). LZ = labyrinth zone. ir = immunoreactive. SpT = spongiotrophoblast. STB = syncytiotrophoblast. NEG CON = negative control. CI = confidence interval.

At GD12.5 in the JZ and LZ, HF diet was associated with decreased ir-PCFT % area stained compared to CON (JZ: B=2.98 [0.73, 5.22], p=0.01; Figure 6A; LZ: B=4.36 [−11.30, 20.02], p=0.05; Figure 6D). However, by GD15.5, HF diet was associated with increased ir-PFCT area stained in the JZ compared to CON (JZ: B=−0.248 [−0.40, −0.09], p=0.004; Figure 6B), but not in the LZ. Neither infection nor HF diet and infection further impacted ir-PCFT % area stained in the JZ or LZ at GD12.5, 15.5 or 18.5 (Figures 6A, B, C, D, E, F).

**Figure 6.**
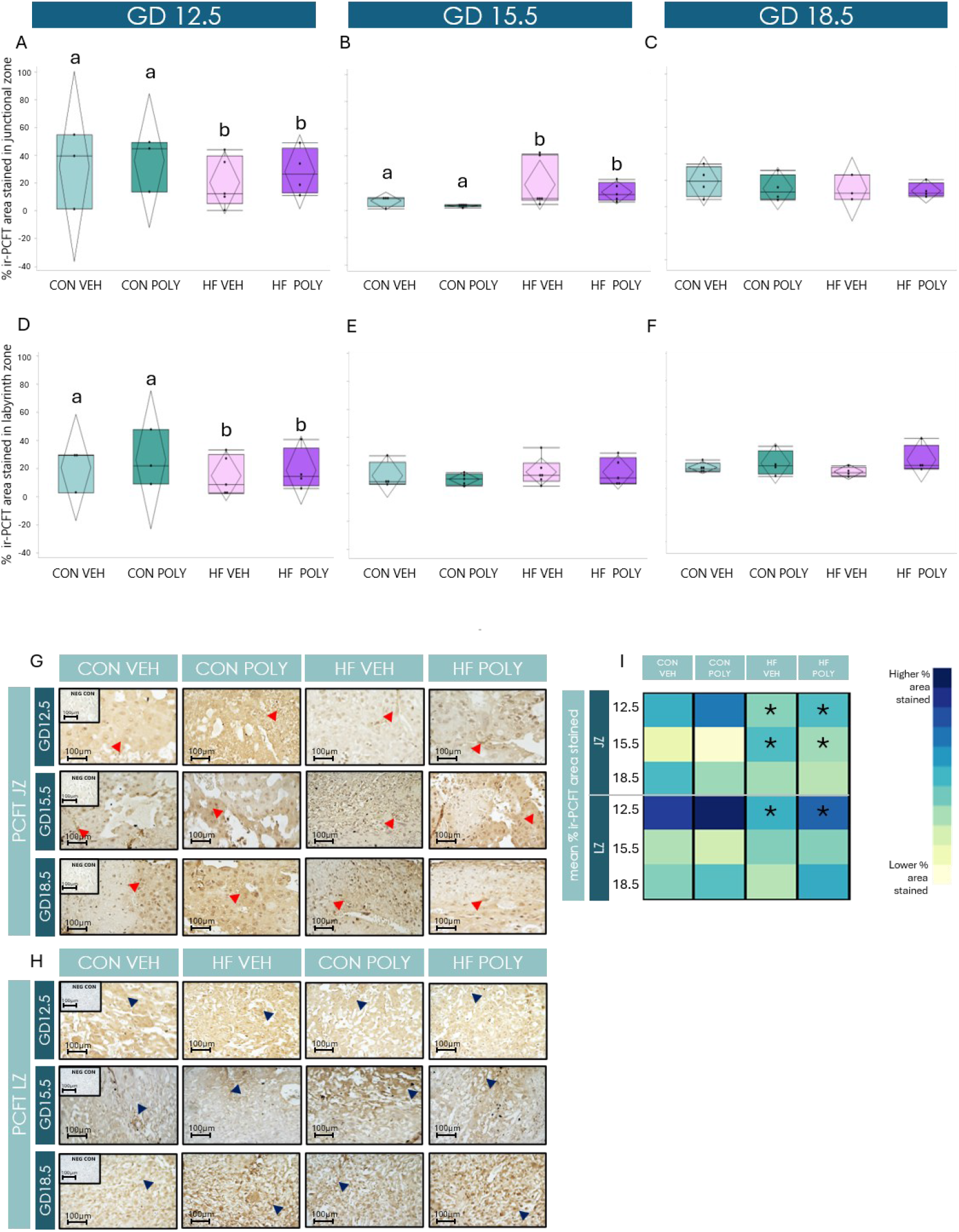
Effect of maternal diet and infectious exposure on placental ir-PCFT % area stained and localisation across gestation. **A, D**. Placental JZ (A) and LZ (D) ir-PCFT % area stained stratified by dietary and infectious exposures at GD12.5. Maternal HF diet was associated with decreased ir-PCFT % area stained in the JZ (p = 0.01) at GD12.5. **B, E**. Placental JZ (B) and LZ (E) ir-PCFT % area stained stratified by dietary and infectious exposures at GD15.5. Maternal HF diet was associated with increased ir-PCFT % area stained in the JZ at GD15.5 compared to CON (JZ: p = 0.004). **C, F**. Placental JZ (C) and LZ (F) ir-PCFT % area stained stratified by dietary and infectious exposures at GD18.5. Data are quantile box plots with 95% CI diamonds. Linear mixed model and Tukey’s post hoc. Groups not connected by the same letter are significantly different, p≤0.05. **G, H**. Representative images of ir-PCFT % area stained in the JZ SpT (G) and LZ STB (H) cytoplasm across dietary and infectious exposures at different GD. Red arrows indicate positive PCFT immunostaining in JZ SpT. Blue arrows indicate positive PCFT immunostaining in LZ STB cytoplasm. Scale bar = 100μm, magnification = 40x. **I**. Heat map of mean % ir-area stained of PCFT in the placental JZ and LZ by GD across exposure groups. Significant differences between dietary and infectious groups in different placental zones and at different GD (p≤0.05) are marked with an asterisk. Darker blue = higher % ir-area stained; Lighter blue/yellow = lower % ir-area stained. CON. GD= gestational day. CON = control. HF = high fat. VEH = vehicle. POLY = Poly(I:C). LZ = labyrinth zone. ir = immunoreactive. SpT = spongiotrophoblast. STB = syncytiotrophoblast. NEG CON = negative control. CI = confidence interval.

Additionally, neither HF diet nor infection impacted placental ir-RFC1 % area stained at any GD (Figure 7F).

**Figure 7.**
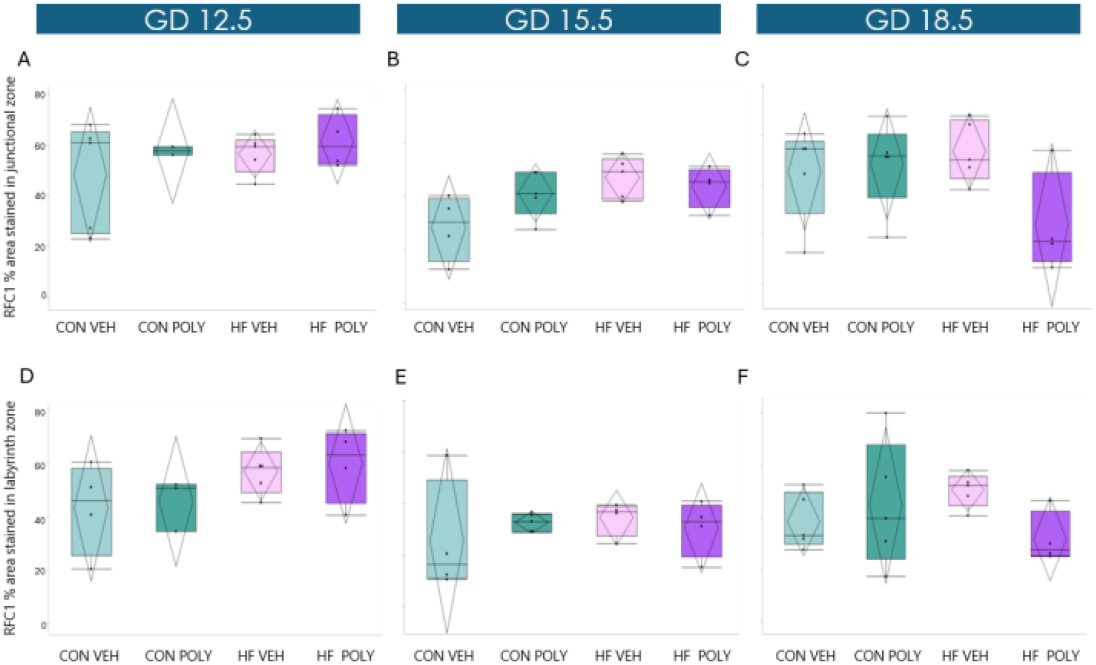
Effect of maternal diet and infectious exposure on placental ir-RFC1 % area stained and localisation across gestation. **A, B, C**. Placental JZ ir-RFC1 % ir-area stained stratified by dietary and infectious exposures at GD12.5, 15.5 and 18.5. **D, E, F**. Placental LZ ir-RFC1 % ir-area stained stratified by dietary and infectious exposures at GD12.5, 15.5 and 18.5. Data are quantile box plots with 95% CI diamonds. Linear mixed model and Tukey’s post hoc. Groups not connected by the same letter are significantly different, p≤0.05. GD = gestational day. ir = immune reactive. CON = control. HF = high fat. VEH = vehicle. POLY = Poly(I:C). JZ = junctional zone LZ = Labyrinth zone. CI = confidence interval.

### Infection, but not HF diet, impacts fetoplacental growth in a gestational-age dependent manner

Next, we evaluated whether and to what extent maternal HF diet or/and infection influenced fetoplacental growth across gestation (Figures 8A, B). Further, we classified placental and fetal weights according to gestational age-specific percentiles as SGA, AGA or LGA to further characterise differences in growth distributions between groups. SGA and LGA represented the lower and upper tails of the placental or fetal weight distributions, respectively, while AGA represented weights within the expected range for gestational age.

**Figure 8.**
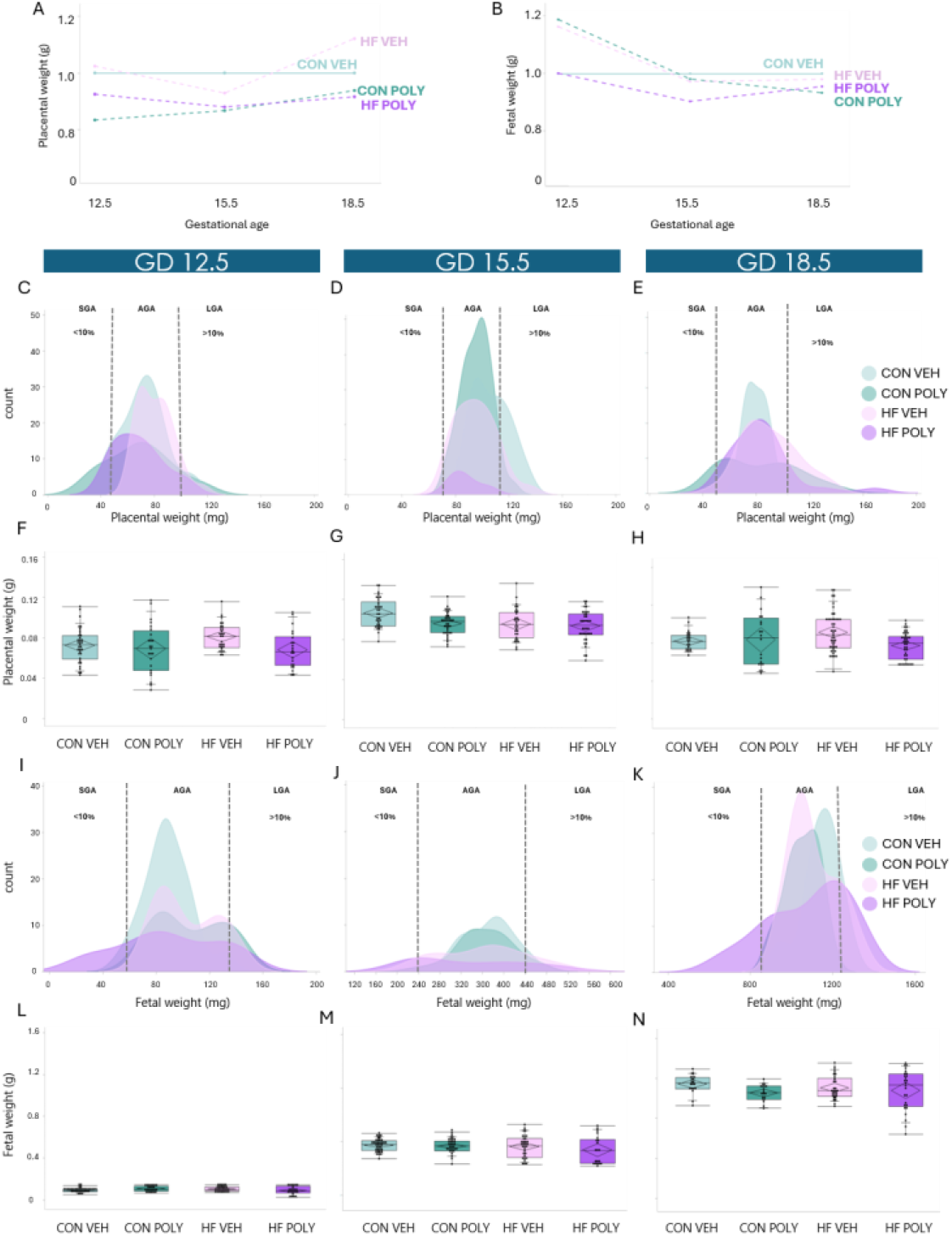
Effect of maternal diet and infectious exposure on placental and fetal weight across gestation. **A, B**. Placental (A) and fetal (B) weight normalised to CON VEH group at GD12.5, 15.5 and 18.5. **C-E**. Distribution of placental weights between dietary and infectious groups across gestation. Kernel density plots show the distribution of placental weights. Vertical dashed lines represent the thresholds defining SGA, <10th percentile, AGA, 10–90th percentile and LGA, >90th percentile. Data are shown for gestational days (C) GD12.5, (D) GD15.5 and (E) GD18.5. Count on the y-axis indicates the number of individual placentae within each weight range. **F-H**. Placental weight stratified by dietary and infectious exposures at (F) GD12.5, (G) GD15.5 and (H) GD18.5. **I-K**. Distribution of fetal weights between dietary and infectious groups across gestation. Kernel density plots show the distribution of fetal weights. Vertical dashed lines represent the thresholds defining SGA, <10th percentile, AGA, 10– 90th percentile and LGA, >90th percentile. Data are shown for gestational days (I) GD12.5, (J) GD15.5, and (K) GD18.5. Count on the y-axis indicates the number of individual fetuses within each weight range. **L-N**. Fetal weight stratified by dietary and infectious exposures at (L) GD12.5, (M) GD15.5 and (N) GD18.5. Data are quantile box plots with 95% CI diamonds. Linear mixed model and Tukey’s post hoc. Groups not connected by the same letter are significantly different, p≤0.05. GD = gestational day. CON = control. HF = high fat. VEH = vehicle. POLY = Poly(I:C). SGA = small for gestational age. AGA = appropriate for gestational age. LGA = large for gestational age. CI = confidence interval.

First, we observed that at GD12.5, a greater proportion of infection-exposed placentae clustered at the lower tail of the placental weight distribution compared to CON VEH (Figure 8C). By GD15.5, placental weight distributions narrowed and groups overlapped, suggesting a more similar growth trajectory in the four groups (Figure 8D). At GD18.5, there was a greater spread in HF VEH, HF POLY and CON POLY placental weights at the lower and upper tails of the placental weight distribution curve (Figure 8E), suggesting that co-exposure to HF diet and infection might be associated with increased variability of placental weight. When quantifying differences in placental weight, we found that neither maternal HF diet nor infection impacted placental weight at any GD (Figure 8A, F-H).

Fetal weight trajectories mirrored the patterns observed in placental growth across gestation. At GD12.5, a greater proportion of HF POLY fetuses skewed towards the lower tail of the fetal weight distribution curve compared to all other groups, and there was a somewhat bimodal distribution of fetal weights for CON POLY and HF VEH (Figure 8I). By GD15.5, HF-exposed fetuses continued to cluster at the lower and upper tail of the distribution compared to CON (Figure 8J). Last, at GD18.5, HF VEH, POLY and CON POLY fetal weight distributions were predominantly right-skewed, with more fetuses in the LGA category, and this was especially true for HF POLY fetuses. HF POLY fetuses were also more represented in the SGA category than all other groups. Collectively these data suggest that co-exposure to HF diet and infection might be associated with increased variability in fetal weight (Figure 8K). When quantifying differences in fetal weight between groups, we found that neither maternal HF diet nor infection impacted fetal weight at any GD (Figures 8B, L-N).

### HF diet and infection associate with sex-dependent differences in placental morphology and function

When stratifying placental data by sex, we observed that male placentae exposed to a HF diet, had increased JZ and LZ area compared to CON at GD18.5 (p≤0.05, Supplementary Figures 1B, D). Sex-specific phenotypes were also observed in placental morphology. For instance, at GD12.5, HF diet was associated with increased maternal blood space area and decreased fetal blood space area compared to CON in male placentae, but not in female placentae (p≤0.05, Supplementary Figure 2B, D). Further, at GD12.5, female placentae exhibited a smaller interhaemal space compared to males (p≤0.05, Supplementary Figure 3A, B). At GD18.5, female placentae exhibited a smaller maternal blood space area compared to males (p≤0.05, Supplementary Figure 3C, D). Additionally, we also observed sex differences in placental folate transporter expression: HF diet was associated with increased *Pcft* mRNA expression compared to CON at GD15.5 in females (p≤0.05, Supplementary Figure 4A), but not males (p≤0.05, Supplementary Figure 4B), and at GD18.5, infectious exposure was associated increased placental *Fr-β* mRNA expression only in male placentae (p≤0.05, Supplementary Figure 4D). Additionally, infection was associated with decreased placental weight in males compared to VEH at GD12.5 (p≤0.05, Supplementary Figure 5B).

## Discussion

We examined placental development and nutrient transporter expression across gestation in pregnancies exposed to maternal HF diet and viral infection. Both exposures independently altered placental structure and nutrient transport. Overall, HF diet disrupted placental development throughout gestation, while at GD12.5, it was associated with increased fatty acid transporter expression and selectively impacted folate transporter expression. HF diet continued to exert a selective effect on folate transporter expression at GD15.5 and decreased folate transporter expression later in gestation. Acute viral infection also modulated placental development in a gestational age-dependent manner, increasing fatty acid transporter expression, followed by decreased folate expression and lower fetal weight later in pregnancy. Although neither HF diet nor infection significantly altered placental or fetal weight at any gestational age, their combined exposure was associated with altered distribution of placental and fetal weights with increased prevalence of SGA and LGA placentae and fetuses throughout gestation. Together, these findings suggest that combined chronic maternal HF diet and acute viral infection may reduce placental resilience leading to maladaptive remodelling and impaired fetoplacental growth.

At GD12.5, maternal HF diet was associated with structural changes in the placenta, which has implications for the growing fetus. We found decreased JZ area size but unaltered placental weight in the face of HF diet, consistent with findings of HF diet and obesity in mouse pregnancy on reduced placental surface area^41^, yet preserved placental weight in later gestation^42^. Decreased JZ area may compromise placental metabolism, impacting its own growth and its capacity to establish an effective nutrient transfer to the fetus^43^. Related, we observed that maternal HF diet was associated with increased maternal blood space area, *Fat/Cd36* mRNA expression and LZ ir-FR-α % area stained, which might suggest placental structural remodelling and functional adaptation to ensure appropriate nutrient supply to the fetus^2^. in the face of HF diet exposure. Additionally, we show for the first time that combined impact of maternal HF diet and infection led to increased JZ ir-FR-α % area stained compared to either exposure alone. This finding suggests that placental folate transport pathways may be particularly sensitive to the combined effects of HF diet and infection challenges, resulting in changes aimed at preserving nutrient availability to the placenta and fetus. Future studies should investigate whether this increase in placental ir-FR-*α* expression results in increased placental folate transport and maintenance of fetal folate status.

In contrast, we observed decreased ir-PCFT % area stained in both JZ and LZ with exposure to HF diet at GD12.5. This is inconsistent with human studies that have not found changes in placental PCFT protein expression in first and second trimester in pregnancies with obesity^32^. Differences between the human data and our findings may relate to methodological differences in where PCFT expression was assessed. In the human study mentioned, PCFT protein expression, along with other folate transporters, was measured in the placental microvillous plasma membrane (MVM)^32^. The MVM is a relevant site for assessing FR-α expression due to its role in direct uptake of folate from the maternal circulation. PCFT, however, is not likely to be localised to the MVM. It is primarily found in the membrane of intracellular endosomes where the microenvironment is ideal for transplacental folate efflux, which will be transferred to the fetal circulation by RFC1^44^. Thus, the lack of PCFT changes reported in human MVM may be due to where PCFT was measured, and not that PCFT expression was truly unchanged. Despite the effect of HF diet in reducing JZ surface area and ir-PCFT % area stained in both JZ and LZ at GD12.5, we did not observe an effect of HF diet on fetal weight, although there were more HF fetuses that were SGA than other groups. This suggests that at GD12.5, the fetus may be somewhat protected from the effects of reduced folate transport capacity.

By GD15.5, exposure to HF diet was associated with, decreased fetal blood space area, and an associated decrease in ir-FR-α in the placental JZ and LZ while an increase ir-PCFT % area stained in the JZ. This is consistent with other studies where obesity in pregnant mice was associated with decreased fetal weight and impaired placental vascularization at GD13.5, changes expected to limit nutrient transfer^45^. Additionally, in human obesity, placental syncytiotrophoblast microvillous plasma membranes (MVM) have been observed to have reduced RFC and FR-α protein expression at term^46^. Collectively these findings suggest that sustained overnutrition during placental development may impair placental growth and nutrient exchange, shown by decreased fetal blood space area, and in our study, may in part explain altered placental folate transport function in the placental JZ. Additionally, it has been previously shown that maternal obesity in mice is associated with placental inflammation at mid-gestation, marked by elevated placental ir-NOD, LRR- and pyrin domain-containing protein 3 (NLRP3, an inflammasome marker) % area stained, and increased placental cell proliferation^29^. Tissue remodelling (evidenced by decreased fetal blood space area) induced by increased inflammasome expression could lead to altered structural organization of the syncytiotrophoblast, which could disrupt the localization or expression of nutrient transporters. These mechanisms could partially explain the reduced FR-α expression observed in HF-exposed placentae. In contrast, the increase in placental ir-PCFT % area stained in the placental JZ at GD15.5 may reflect a compensatory response to maintain folate supply to the fetus under HF diet exposure. Despite these changes observed in decreased fetal blood space area and select folate transport, fetal weight was unaffected by HF and/or infection at this stage of pregnancy, although there was a greater distribution of weights from SGA through to LGA in HF fetuses compared to CON. These data suggest that select nutrient transporters may adapt to temporarily preserve fetal growth in mid-pregnancy.

By GD18.5 HF diet was associated with increased JZ and LZ area size, but placental weights were not altered. Given the exponential fetal growth towards the end of normal pregnancy, the observed change from reduced placental morphometric features at GD12.5 and 15.5 to increased in late pregnancy may reflect placental adaptations to support an appropriate fetal growth trajectory in the face of nutritional challenge. Indeed, we saw an increase in the number of AGA and LGA fetuses by end of pregnancy in HF groups. These findings also align with our previous work, where we found that maternal HF diet was associated with increased placental LZ cell proliferation at GD15.5^29^, which may precede the observed increases in placental zone areas we observed in the present study. Additionally in late pregnancy, folate transport continued to be impacted by HF diet, where we observed decreased ir-Fr-α % area stained in the JZ. This zone-specific reduction could limit folate-dependent endocrine and growth-regulatory functions of the JZ with implications for fetal development.

Despite limited research investigating how infection influences placental morphology and nutrient transporter expression throughout pregnancy, our study provides novel evidence that acute viral infection dynamically modulates placental development and fatty acid and folate transport across gestation. For instance, we found that infection was associated with increased *Fat/Cd36* mRNA expression at GD12.5. CD36 also functions as a signaling receptor responding to pathogen-associated molecular patterns (PAMPs), which are elevated during viral infectious exposure^47^. This upregulation may lead to increased nutrient transfer to the fetus^47^. However, despite that some changes reported in the current study suggest placental adaptations to support optimal fetoplacental growth, infection was associated with decreased placental weight at GD12.5 and 15.5, suggesting some degree of placental insufficiency with acute viral infection. Infectious exposure also had an important effect at later pregnancy where we show for the first time decreased *Rfc1* mRNA expression, in response to viral infection, supporting the notion that the placenta failed to adequately compensate for its altered development and function across pregnancy in the face of a second challenge. These findings highlight the placenta’s limited plasticity when subjected to concurrent stressors. Maternal chronic stress perhaps induced by obesity/prolonged HF diet, could lead to placental exhaustion, reducing the placenta’s capacity to respond and adapt effectively to additional insults such as viral infection^48,49^.

Last, we explored whether and to what extent changes in placental development in response to maternal HF diet and acute viral infection were different between male and female placentae. Overall, male placentae exhibited greater growth adaptations, whereas female placentae showed more pronounced alterations in placental architecture at the maternal-fetal exchange interface and nutrient transporter expression. Studies have shown that male placentae prioritize intrauterine growth, often maintaining growth trajectories despite suboptimal conditions, compared to females *in utero*^*50*^. In particular, pregnancies carrying female fetuses have been associated with altered placental growth trajectories and activation of stress and inflammatory response pathways, which could influence placental architecture and expression and localisation of nutrient transporters^51^. Indeed, we found that HF diet at GD15.5 was associated with increased LZ area compared to CON in males, but not females. Further, male placentae exhibited an increased maternal blood space area at GD12.5, while concomitant decreased in fetal blood space area compared to CON. These findings may reflect a response to maintain nutrient supply to the growing fetus in males at GD12.5. Although, female placentae exhibited a smaller interhaemal space at GD12.5, at GD18.5 females showed a smaller maternal blood space area compared to males. Additionally, in females, we observed increased *Pcft* mRNA expression in response to HF diet. Additionally, at GD18.5 we found that infection was associated with increased *Fr-β* mRNA expression in males but not females. Infectious exposure also impacted placental growth in a sex-specific manner: acute infection was associated with decreased placental weight at GD12.5 in males, but not females. These data suggest distinct sex-specific adaptations to maternal diet and infection, which may depend on the developmental window. Data from normal pregnancies show that male fetuses are generally heavier at birth than females^52^. Our results suggest that in pregnancies exposed to HF diet and infection, although male placentae exhibit structural adaptations that may optimize maternal-fetal exchange, they remain susceptible to decreased placental weight.

Our study has several strengths. First, we provide novel insight into how placental fatty acid and folate transporters respond to metabolic and immune stressors across key developmental stages in pregnancy, which has not been previously done. Additionally, our dietary model controlled for fibre-associated effects on the maternal gut and systemic inflammation^53^ and mirror human dietary patterns, which, are not as fibre-restricted as most experimental HF diets^54,55^, as we described previously^29^. Reduced dietary fibre can impair intestinal nutrient absorption and lower maternal circulating nutrient availability, potentially confounding interpretation of placental nutrient transporter expression^56^. Further, our Poly(I:C) model provided a controlled, acute viral-like challenge, enabling precise assessment of how infection timing influences placental development and function^57,58^. Finally, our study also explored sex differences in placental development and function across gestation, an aspect that has been relatively underexplored in developmental research. Nonetheless, some limitations to our work exist. Although select nutrient transporter expression was characterised, direct functional assessments of nutrient transfer were not assessed, leaving it unclear whether the observed alterations in transporter expression result in altered nutrient availability for the fetus. Additionally, sample sizes for certain histological analyses were limited to three placentae per group/GD and should be interpreted cautiously. Replication studies with large sample sizes will be important to confirm the robustness of these findings. Last, while we explored sex differences in the expression of nutrient transporters and fetoplacental development, future studies should clarify the mechanisms underlying these sex-specific placental responses to maternal HF diet and infection and assess their contribution to sex differences in offspring growth and long-term health.

We found that maternal HF diet and viral infection independently disrupt placental structure and nutrient transport, in a gestational age-dependent manner. Maternal HF diet induced sustained alterations in placental morphology, fatty acid transporter and folate expression. Importantly, the combined exposure of maternal HF diet and infection increased folate transporter expression beyond that observed with either exposure alone, suggesting that placental folate transport mechanisms may be particularly sensitive and responsive to co-exposed HF and infectious stress. Together, our data emphasise the placenta’s central role as both a target and mediator of maternal HF diet and infectious challenges, reinforcing the need for strategies that mitigate maternal overnutrition and viral infection to improve pregnancy outcomes.

## Data availability

Some or all datasets generated during and/or analysed during the current study are available from the corresponding author on reasonable request.

## Author Contributions

Conceptualisation, methodology: TF, SM, LC, EB, KLC; formal analysis: TF, LC; writing—original draft preparation: TF, KLC; writing—review and editing: TF, LC, EB, KLC; funding acquisition, resources, supervision: KLC.

## Acknowledgments

We would like to acknowledge the funding from the Natural Sciences and Engineering Research Council of Canada (NSERC; RGPIN-2018-05300 and DGECR-2018-00203). Hailey Scott (Department of Health Sciences, Carleton University) for her contribution and assistance throughout the execution and write up of this project, especially in the qPCR methodology and peer review. Erikles Barbosa (Department of Health Sciences, Carleton University) for his contribution during peer review. Israa Zareef, Oluwatomike Aribaloye and Tiffany Kerdilès (Department of Health Sciences, Carleton University) for their assistance during sample collection and pilot study execution.

## Supplementary tables

**Supplementary Table 1.**
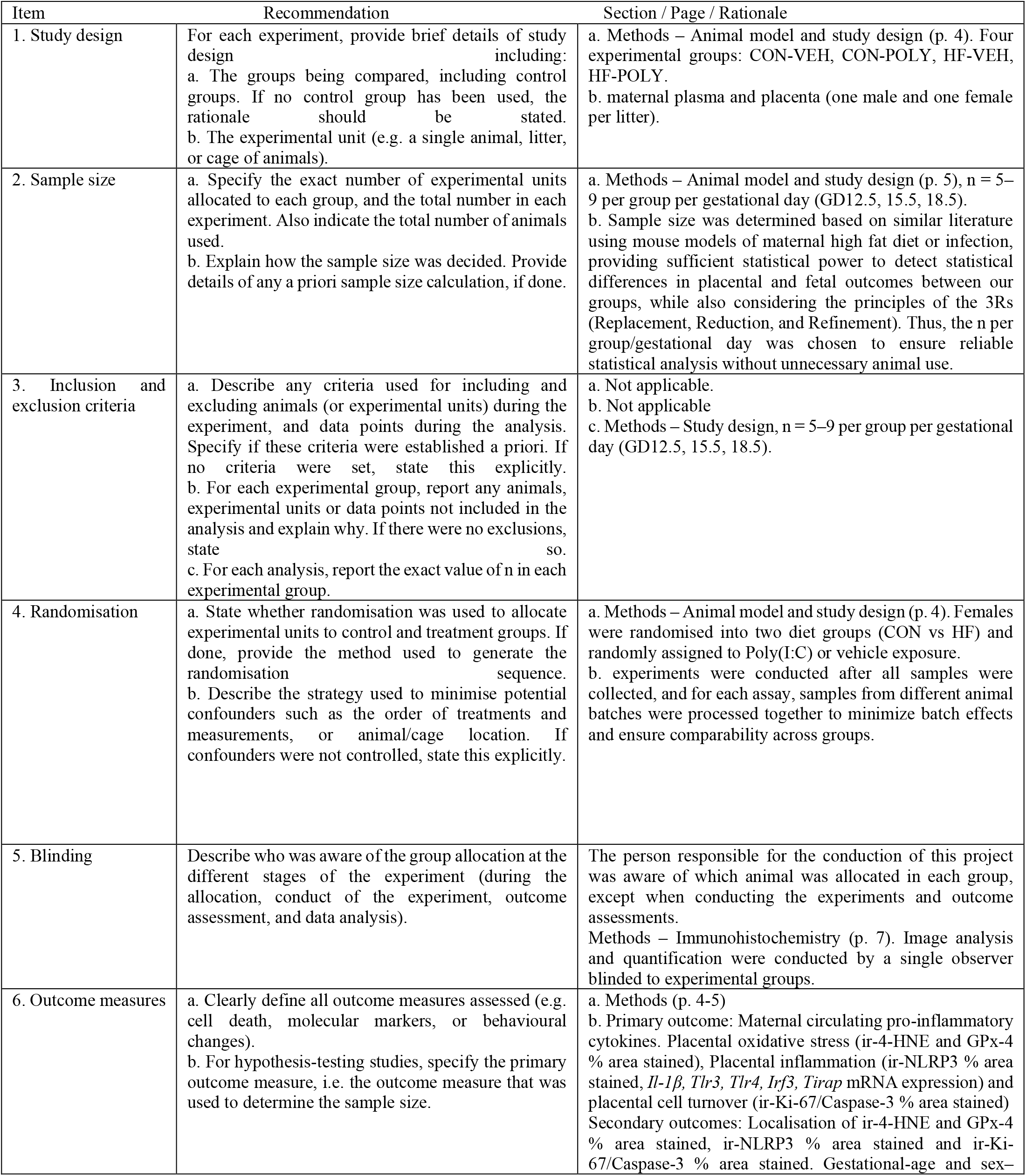
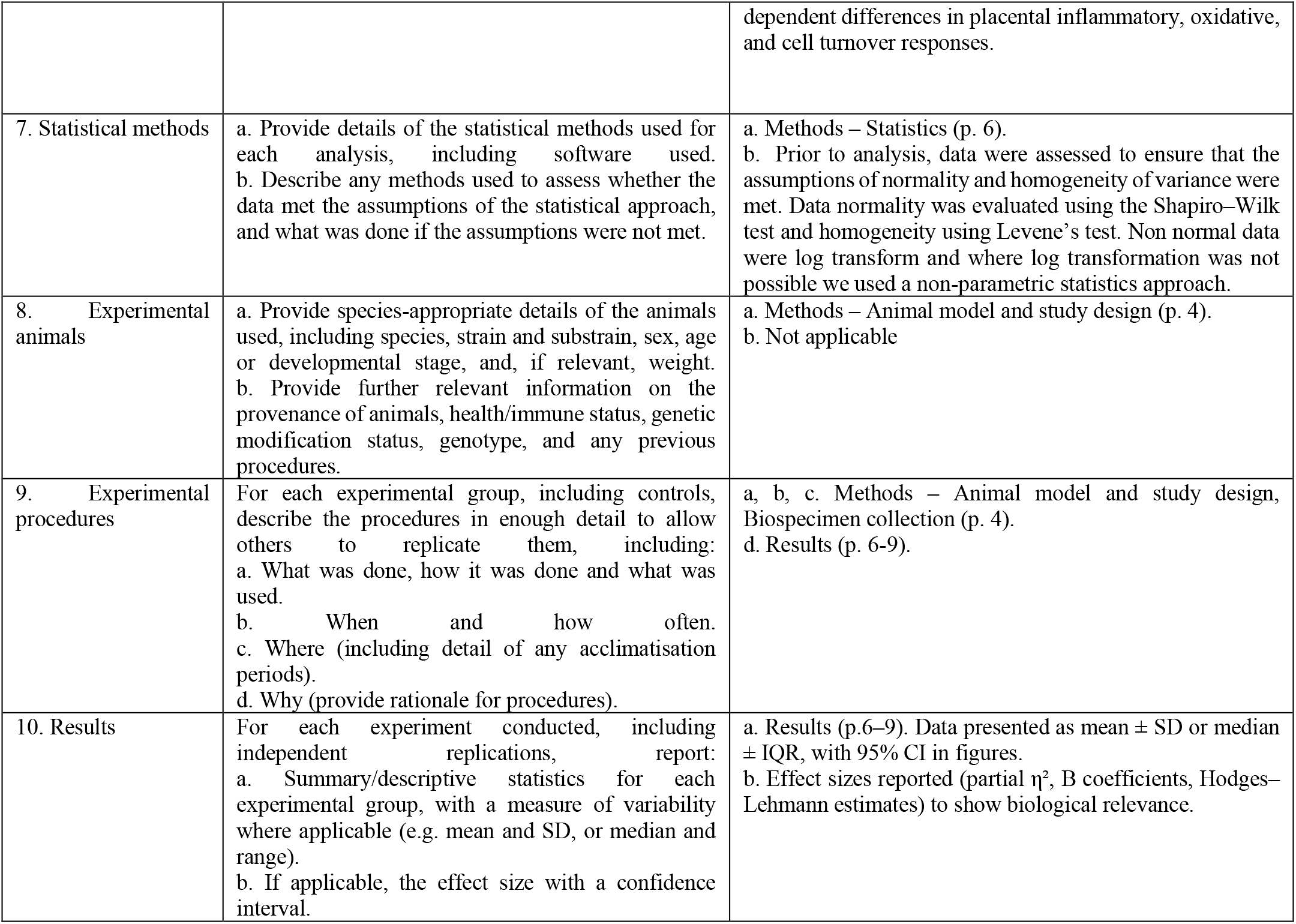
ARRIVE Essential 10 – Compliance Summary. These items are the basic minimum to include in a manuscript. Without this information, readers and reviewers cannot assess the reliability of the findings.

**Supplementary Table 2.**
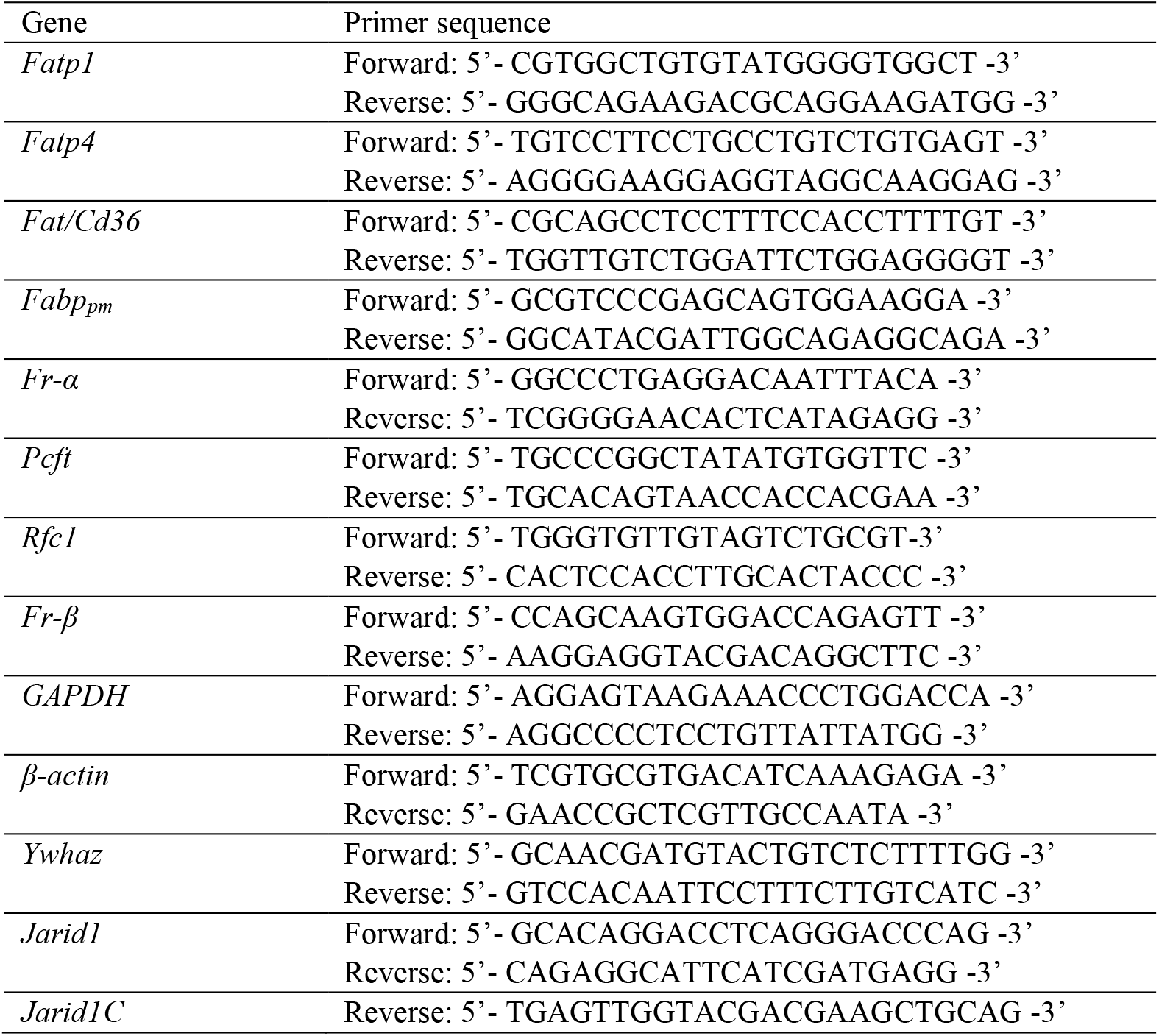
Primer sequences for real-time quantitative PCR gene studies.

## Supplementary Figures

**Supplementary Figure 1.**
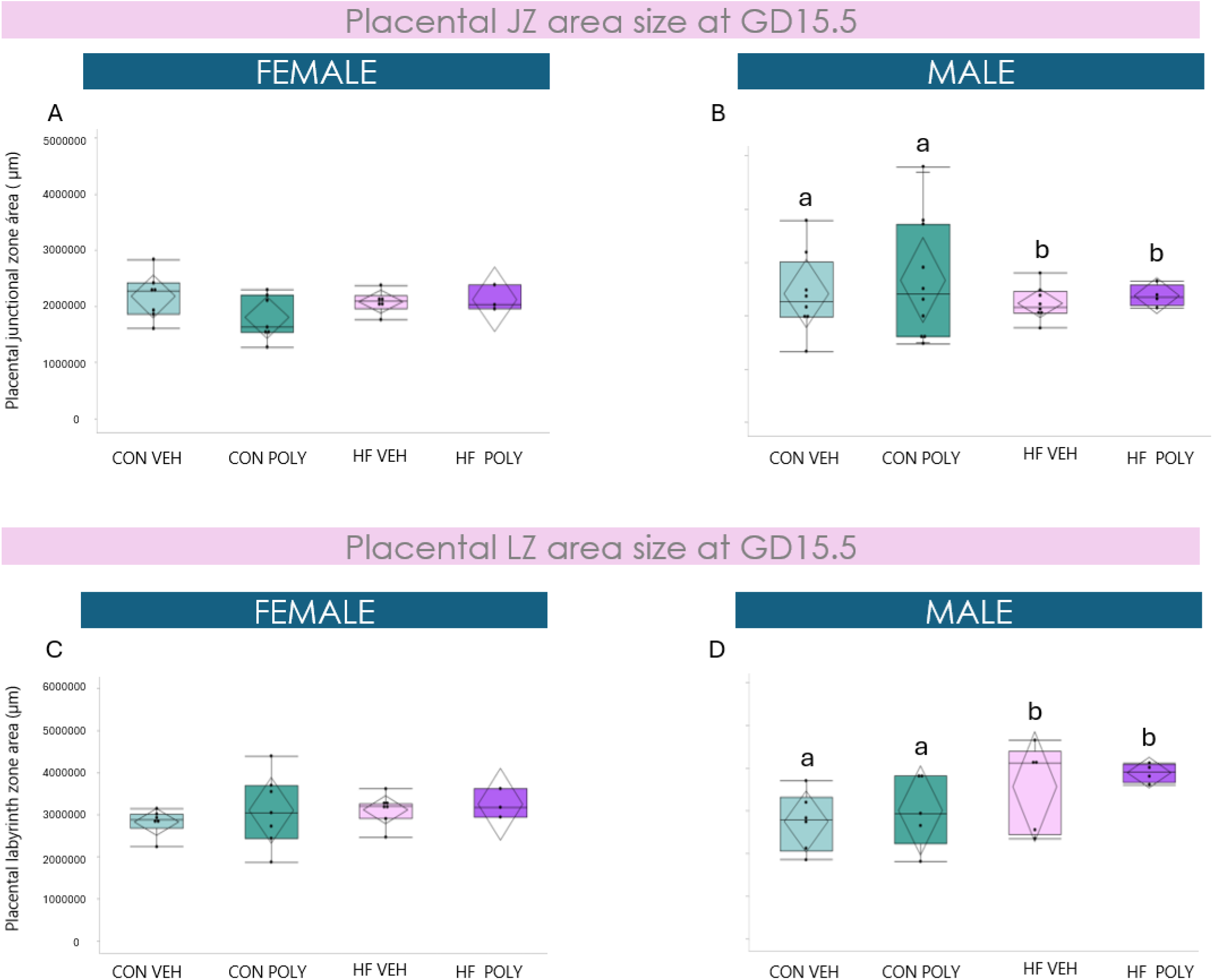
Effect of maternal HF diet and infectious exposure on placental JZ and LZ area stratified by sex at GD15.5. **A**. Placental JZ area of female fetuses stratified by diet and infectious exposure groups at GD15.5. **B**. Placental JZ area size of male fetuses stratified by diet and infectious exposure groups at GD15.5. Maternal HF diet exposure associated with decreased placental JZ area compared to CON (p=0.03). **C**. Placental LZ area of female fetuses stratified by diet and infectious exposure groups at GD15.5. **D**. Placental LZ area of male fetuses stratified by diet and infectious exposure groups at GD15.5. Maternal HF diet exposure associated with increased placental LZ area compared to CON (p=0.02). Data are quantile box plots with 95% CI diamonds. Two-way ANOVA and Tukey’s post hoc, p≤0.05. Groups not connected by the same letter are significantly different. GD = gestational day. CON = control. HF = high fat. VEH = vehicle. POLY = Poly(I:C). LZ = labyrinth zone. JZ = junctional zone.

**Supplementary Figure 2.**
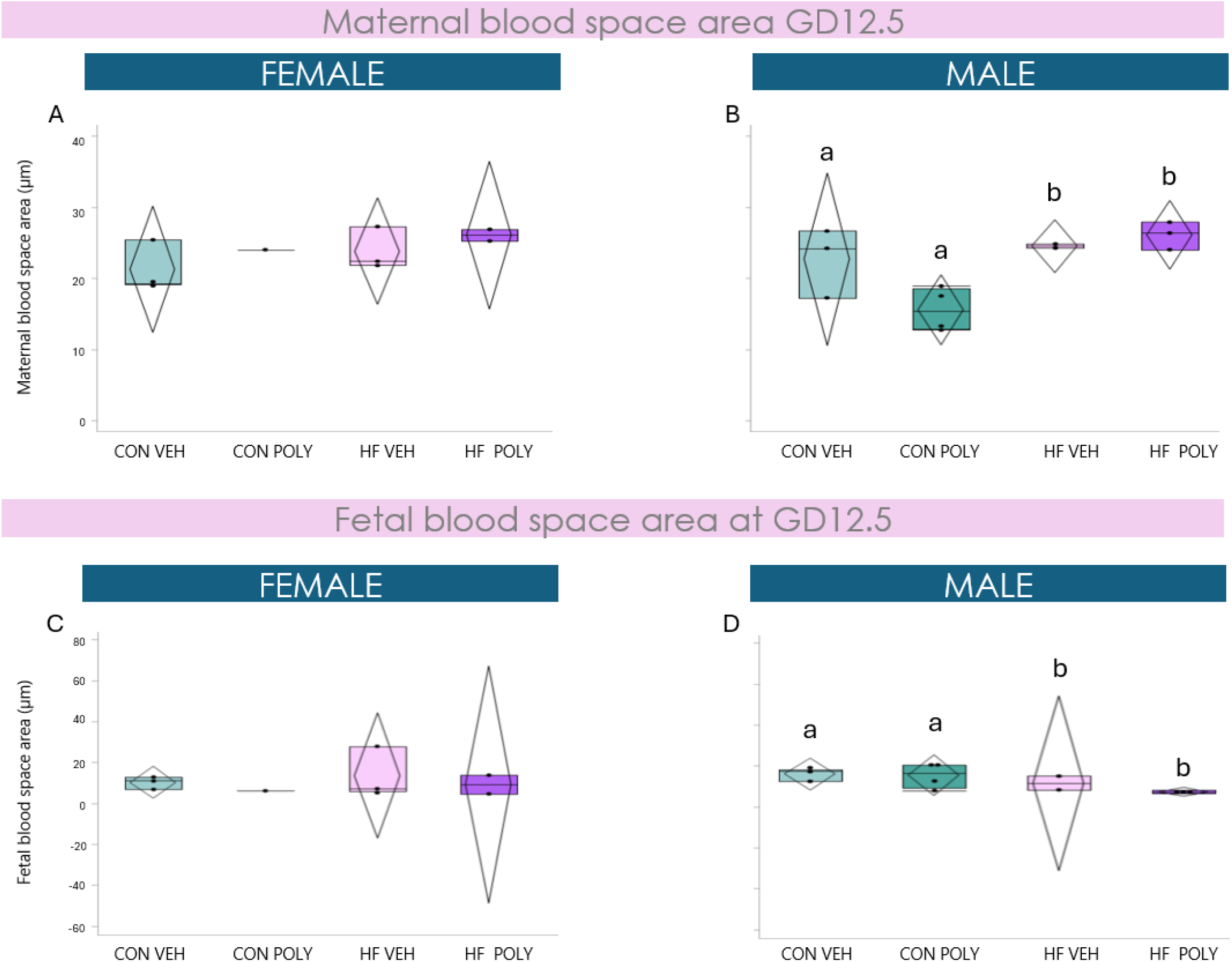
Effect of maternal HF diet and infectious exposure on placental blood space area stratified by sex across pregnancy. **A, B**. Maternal blood space area at GD12.5 stratified by dietary and infectious exposures across gestation in female and male placentae. **B**. Maternal HF diet associated with increased maternal blood space area compared to CON (p=0.01) in male placentae. **C, D**. Fetal blood space area at GD12.5 stratified by dietary and infectious exposures across gestation in female and male placentae. **D**. Maternal HF diet associated with decreased fetal blood space area compared to CON (p=0.03) in male placentae. Data are quantile box plots with 95% CI diamonds. Two-way ANOVA and Tukey’s post hoc, p≤0.05. Groups not connected by the same letter are significantly different. GD = gestational day. CON = control. HF = high fat. VEH = vehicle. POLY = Poly(I:C).

**Supplementary Figure 3.**
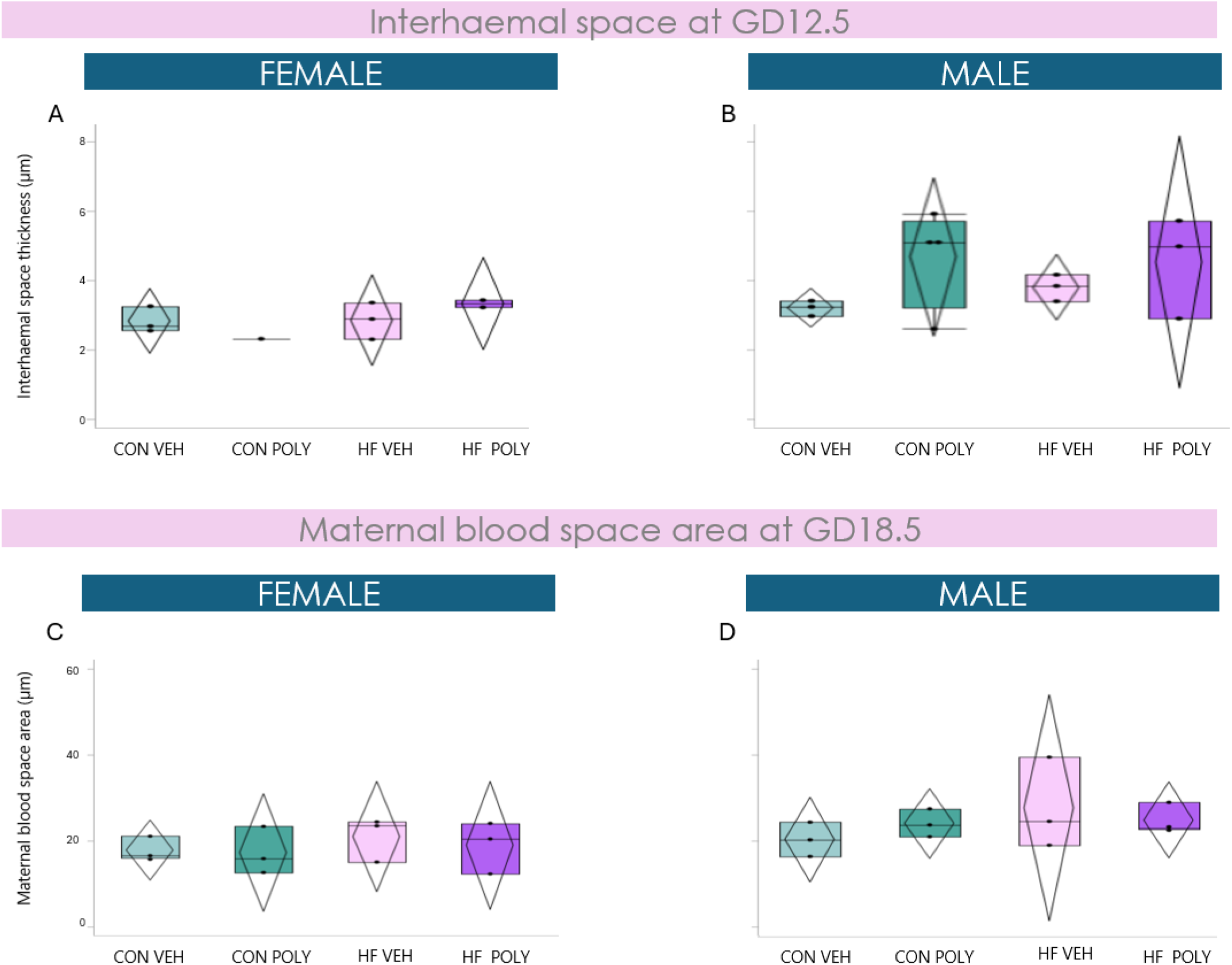
Effect of maternal HF diet and infectious exposure on placental interhaemal space and blood space stratified by sex across pregnancy. **A, B**. Placental interhaemal space stratified by dietary and infectious exposures across gestation in female and male placentae. At GD12.5, female placentae exhibited a smaller interhaemal space compared to males (p=0.03). **E, F**. Maternal blood space are at GD18.5 stratified by dietary and infectious exposures across gestation in female and male placentae. At GD18.5, male placentae exhibited a greater maternal blood space area compared to females (p=0.02). Data are quantile box plots with 95% CI diamonds. Three-way ANOVA and Tukey’s post hoc, p≤0.05. Groups not connected by the same letter are significantly different. GD = gestational day. CON = control. HF = high fat. VEH = vehicle. POLY = Poly(I:C).

**Supplementary Figure 4.**
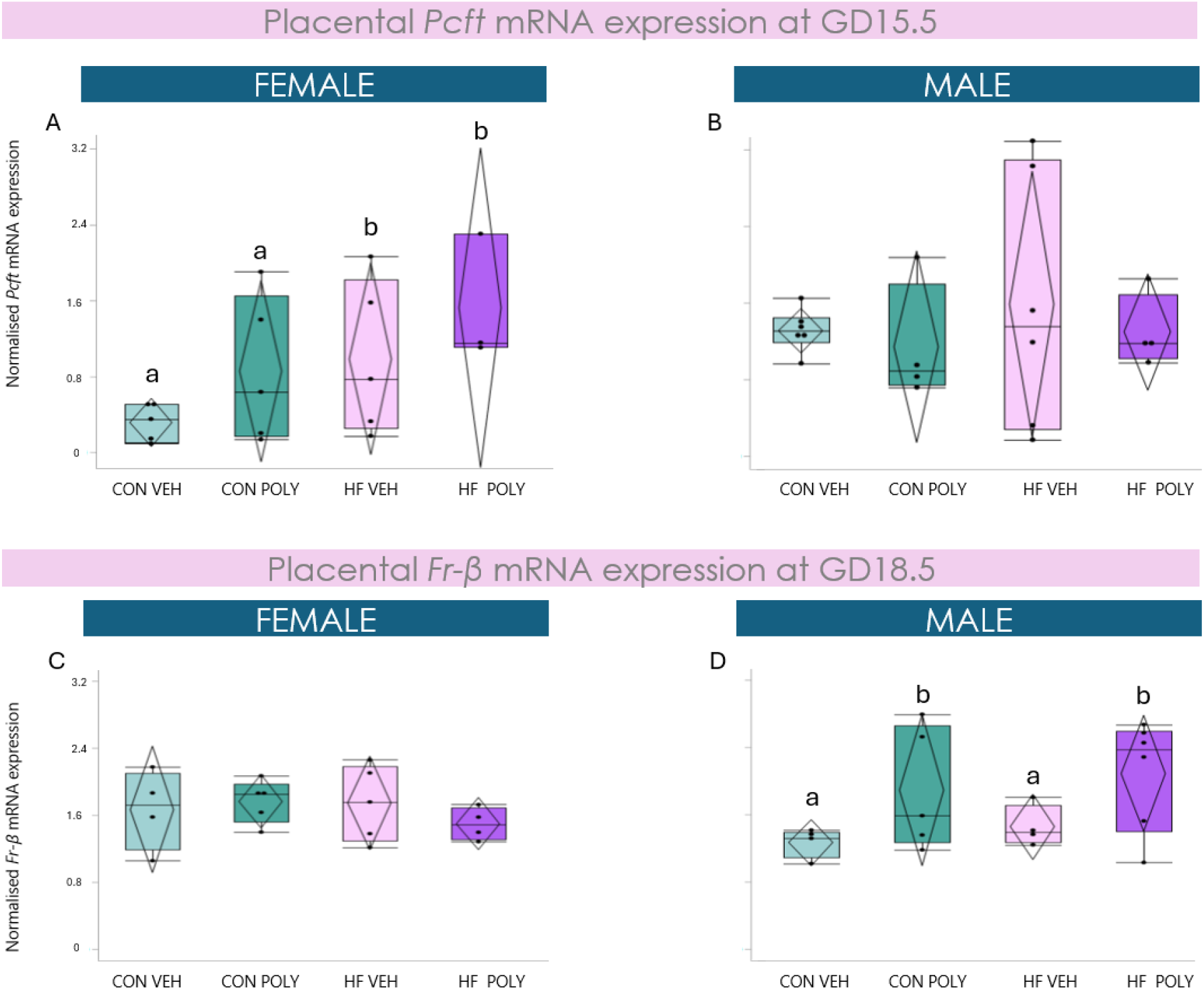
Effect of maternal HF diet and infectious exposure on placental folate transporter mRNA expression stratified by sex at GD15.5 and 18.5. **A**. Placental *Pcft* mRNA expression in female fetuses stratified by diet and infectious exposure groups at GD12.5. Maternal HF diet exposure associated with increased *Pcft* mRNA expression compared to CON (p=0.04). **B**. Placental *Pcft* mRNA expression in male fetuses stratified by diet and infectious exposure groups at GD12.5. **C**. Placental *Fr-β* mRNA expression in female fetuses stratified by diet and infectious exposure groups at GD18.5. **D**. Placental *Fr-β* mRNA expression in male fetuses stratified by diet and infectious exposure groups at GD18.5. Poly(I:C) exposure associated with increased *Fr-β* mRNA expression compared to VEH (p=0.03). Data are quantile box plots with 95% CI diamonds. Two-way ANOVA and Tukey’s post hoc, p≤0.05. Groups not connected by the same letter are significantly different. GD = gestational day. CON = control. HF = high fat. VEH = vehicle. POLY = Poly(I:C).

**Supplementary Figure 5.**
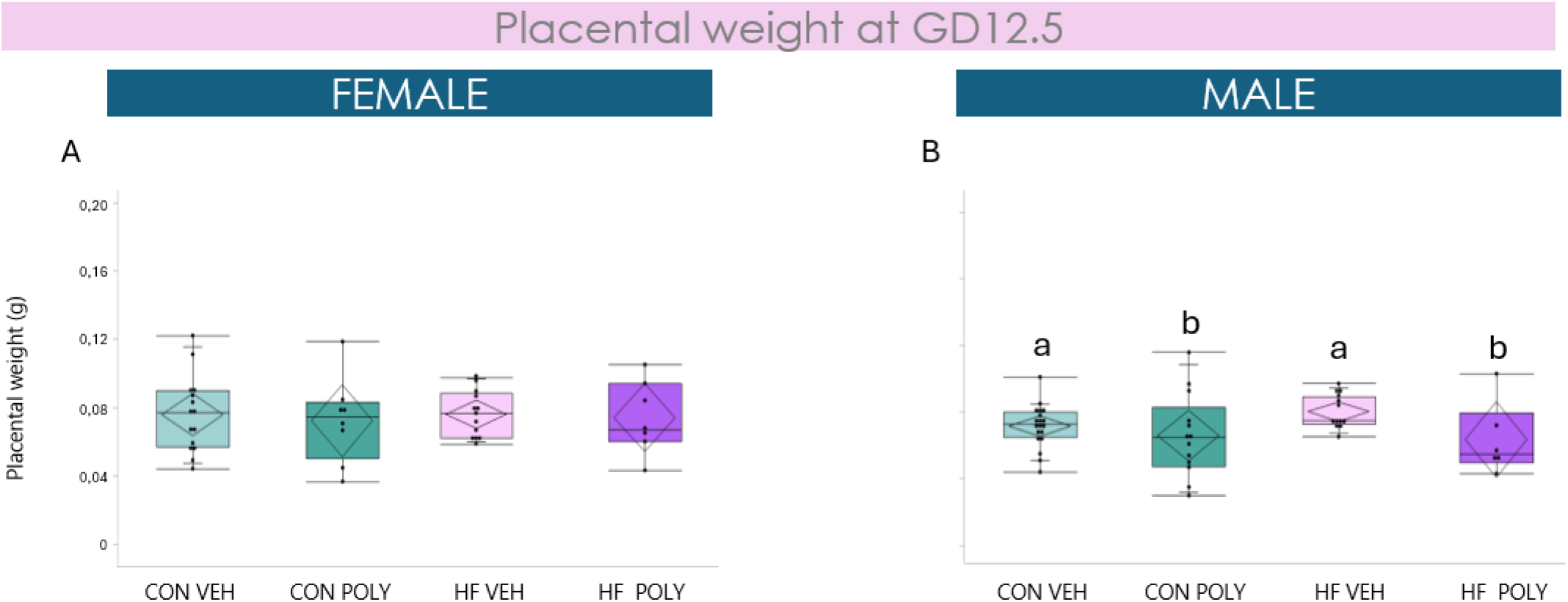
Effect of maternal HF diet and infectious exposure on placental weight stratified by sex at GD12.5. **A**. Placental weight of female fetuses stratified by diet and infectious exposure groups at GD12.5. **B**. Placental weight of male fetuses stratified by diet and infectious exposure groups at GD12.5. Poly(I:C) exposure associated with decreased placental weight compared to CON (p=0.02). Data are quantile box plots with 95% CI diamonds. Two-way ANOVA and Tukey’s post hoc, p≤0.05. Groups not connected by the same letter are significantly different. GD = gestational day. CON = control. HF = high fat. VEH = vehicle. POLY = Poly(I:C).

## References

1. Brett KE, Ferraro ZM, Yockell-Lelievre J, Gruslin A, Adamo KB. Maternal-fetal nutrient transport in pregnancy pathologies: the role of the placenta. Int J Mol Sci. Sep 12 2014;15(9):16153–85. doi:10.3390/ijms150916153

2. Connor KL, Kibschull M, Matysiak-Zablocki E, et al. Maternal malnutrition impacts placental morphology and transporter expression: an origin for poor offspring growth. J Nutr Biochem. Jan 8 2020;78:108329. doi:10.1016/j.jnutbio.2019.108329

3. Abu-Raya B, Michalski C, Sadarangani M, Lavoie PM. Maternal Immunological Adaptation During Normal Pregnancy. Front Immunol. 2020;11:575197. doi:10.3389/fimmu.2020.575197

4. Burton GJ, Fowden AL, Thornburg KL. Placental Origins of Chronic Disease. Physiol Rev. 10 2016;96(4):1509–65. doi:10.1152/physrev.00029.2015

5. Ogden CL, Carroll MD, Fryar CD, Flegal KM. Prevalence of Obesity Among Adults and Youth: United States, 2011-2014. NCHS Data Brief. Nov 2015;(219):1–8.

6. McClymont E, Albert AY, Alton GD, et al. Association of SARS-CoV-2 Infection During Pregnancy With Maternal and Perinatal Outcomes. JAMA. May 24 2022;327(20):1983–1991. doi:10.1001/jama.2022.5906

7. Karlsson EA, Marcelin G, Webby RJ, Schultz-Cherry S. Review on the impact of pregnancy and obesity on influenza virus infection. Influenza Other Respir Viruses. Nov 2012;6(6):449–60. doi:10.1111/j.1750-2659.2012.00342.x

8. Creisher PS, Perry JL, Zhong W, et al. Adverse outcomes in SARS-CoV-2-infected pregnant mice are gestational age-dependent and resolve with antiviral treatment. J Clin Invest. Oct 16 2023;133(20)doi:10.1172/JCI170687

9. Tan L, Lacko LA, Zhou T, et al. Pre- and peri-implantation Zika virus infection impairs fetal development by targeting trophectoderm cells. Nat Commun. Sep 13 2019;10(1):4155. doi:10.1038/s41467-019-12063-2

10. Wang R, Yan W, Du M, Tao L, Liu J. The effect of influenza virus infection on pregnancy outcomes: A systematic review and meta-analysis of cohort studies. Int J Infect Dis. Apr 2021;105:567–578. doi:10.1016/j.ijid.2021.02.095

11. Cherukad J, Wainwright V, Watson ED. Spatial and temporal expression of folate-related transporters and metabolic enzymes during mouse placental development. Placenta. May 2012;33(5):440–8. doi:10.1016/j.placenta.2012.02.005

12. Islam A, Kagawa Y, Sharifi K, et al. Fatty Acid Binding Protein 3 Is Involved in n-3 and n-6 PUFA transport in mouse trophoblasts. J Nutr. Oct 2014;144(10):1509–16. doi:10.3945/jn.114.197202

13. Duttaroy AK, Basak S. Maternal Fatty Acid Metabolism in Pregnancy and Its Consequences in the Feto-Placental Development. Front Physiol. 2021;12:787848. doi:10.3389/fphys.2021.787848

14. Kalhan TG, Bateman DA, Bowker RM, Hod EA, Kashyap S. Effect of red blood cell storage time on markers of hemolysis and inflammation in transfused very low birth weight infants. Pediatr Res. Dec 2017;82(6):964–969. doi:10.1038/pr.2017.177

15. Zhang XM, Huang GW, Tian ZH, Ren DL, Wilson JX. Folate stimulates ERK1/2 phosphorylation and cell proliferation in fetal neural stem cells. Nutr Neurosci. Oct 2009;12(5):226–32. doi:10.1179/147683009×423418

16. Greenberg JA, Bell SJ, Guan Y, Yu YH. Folic Acid supplementation and pregnancy: more than just neural tube defect prevention. Rev Obstet Gynecol. Summer 2011;4(2):52–9.

17. de Souza Lima B, Sanches APV, Ferreira MS, de Oliveira JL, Cleal JK, Ignacio-Souza L. Maternal-placental axis and its impact on fetal outcomes, metabolism, and development. Biochim Biophys Acta Mol Basis Dis. Jan 2024;1870(1):166855. doi:10.1016/j.bbadis.2023.166855

18. Huang Z, Huang S, Song T, Yin Y, Tan C. Placental Angiogenesis in Mammals: A Review of the Regulatory Effects of Signaling Pathways and Functional Nutrients. Adv Nutr. Dec 01 2021;12(6):2415–2434. doi:10.1093/advances/nmab070

19. Dumolt JH, Powell TL, Jansson T. Placental Function and the Development of Fetal Overgrowth and Fetal Growth Restriction. Obstet Gynecol Clin North Am. Jun 2021;48(2):247–266. doi:10.1016/j.ogc.2021.02.001

20. Rosario FJ, Nathanielsz PW, Powell TL, Jansson T. Maternal folate deficiency causes inhibition of mTOR signaling, down-regulation of placental amino acid transporters and fetal growth restriction in mice. Sci Rep. Jun 21 2017;7(1):3982. doi:10.1038/s41598-017-03888-2

21. Kolahi KS, Valent AM, Thornburg KL. Real-time microscopic assessment of fatty acid uptake kinetics in the human term placenta. Placenta. Dec 2018;72-73:1–9. doi:10.1016/j.placenta.2018.07.014

22. Johnsen GM, Basak S, Weedon-Fekjær MS, Staff AC, Duttaroy AK. Docosahexaenoic acid stimulates tube formation in first trimester trophoblast cells, HTR8/SVneo. Placenta. Sep 2011;32(9):626–632. doi:10.1016/j.placenta.2011.06.009

23. Liu L, Zhao H, Wang Y, et al. Docosahexaenoic acid insufficiency impairs placental angiogenesis by repressing the methylene-bridge fatty acylation of AKT in preeclampsia. Placenta. Sep 26 2024;155:100–112. doi:10.1016/j.placenta.2024.08.012

24. Delhaes F, Giza SA, Koreman T, et al. Altered maternal and placental lipid metabolism and fetal fat development in obesity: Current knowledge and advances in non-invasive assessment. Placenta. Sep 2018;69:118–124. doi:10.1016/j.placenta.2018.05.011

25. Isabel C, Faro Rebecca V, Vrijkotte TGM, Theodorus Bartholomeus T. Early pregnancy triglycerides and not fructosamine are associated with birth weight (with foetal sexual dimorphism). Eur J Clin Invest. Jan 2023;53(1):e13896. doi:10.1111/eci.13896

26. Duttaroy AK, Basak S. Maternal dietary fatty acids and their roles in human placental development. Prostaglandins Leukot Essent Fatty Acids. Apr 2020;155:102080. doi:10.1016/j.plefa.2020.102080

27. Dubé E, Gravel A, Martin C, et al. Modulation of fatty acid transport and metabolism by maternal obesity in the human full-term placenta. Biol Reprod. Jul 2012;87(1):14, 1-11. doi:10.1095/biolreprod.111.098095

28. van Vliet MM, Schoenmakers S, Willemsen SP, Sinclair KD, Steegers-Theunissen RPM. First-trimester maternal folate and vitamin B12 concentrations and their associations with first-trimester placental growth: the Rotterdam Periconception Cohort. Hum Reprod. Aug 01 2025;40(8):1485–1494. doi:10.1093/humrep/deaf095

29. Ferraz T, Cardoso L, Mohammadkhani S, Bloise E, Connor KL. Maternal High Fat Diet and Acute Viral Mimic Exposure Impact Placental Inflammation, Lipid Peroxidation and Cellular Proliferation-to-Death Ratio Across Mouse Pregnancy. Am J Reprod Immunol. Aug 2026;96(2):e70290. doi:10.1111/aji.70290

30. Cervantes EM, Girard S. Placental Inflammation in Preterm Premature Rupture of Membranes and Risk of Neurodevelopmental Disorders. Cells. Jun 24 2025;14(13)doi:10.3390/cells14130965

31. Aye IL, Lager S, Ramirez VI, et al. Increasing maternal body mass index is associated with systemic inflammation in the mother and the activation of distinct placental inflammatory pathways. Biol Reprod. Jun 2014;90(6):129. doi:10.1095/biolreprod.113.116186

32. Jessel RH, Rosario FJ, Chen YY, et al. Decreased placental folate transporter expression and activity in first and second trimester in obese mothers. J Nutr Biochem. Mar 2020;77:108305. doi:10.1016/j.jnutbio.2019.108305

33. Palladino E, Van Mieghem T, Connor KL. Diet Alters Micronutrient Pathways in the Gut and Placenta that Regulate Fetal Growth and Development in Pregnant Mice. Reprod Sci. 02 2021;28(2):447–461. doi:10.1007/s43032-020-00297-1

34. Hsia JZ, Liu D, Haynes L, Cruz-Cosme R, Tang Q. Lipid Droplets: Formation, Degradation, and Their Role in Cellular Responses to Flavivirus Infections. Microorganisms. Mar 24 2024;12(4)doi:10.3390/microorganisms12040647

35. Pfeifer E, Parrott J, Lee GT, et al. Regulation of human placental drug transporters in HCV infection and their influence on direct acting antiviral medications. Placenta. Sep 2018;69:32–39. doi:10.1016/j.placenta.2018.07.005

36. Percie du Sert N, Hurst V, Ahluwalia A, et al. The ARRIVE guidelines 2.0: Updated guidelines for reporting animal research. PLoS Biol. Jul 2020;18(7):e3000410. doi:10.1371/journal.pbio.3000410

37. Hemberger M, Hanna CW, Dean W. Mechanisms of early placental development in mouse and humans. Nat Rev Genet. Jan 2020;21(1):27–43. doi:10.1038/s41576-019-0169-4

38. Elmore SA, Cochran RZ, Bolon B, et al. Histology Atlas of the Developing Mouse Placenta. Toxicol Pathol. Jan 2022;50(1):60–117. doi:10.1177/01926233211042270

39. Livak KJ, Schmittgen TD. Analysis of relative gene expression data using real-time quantitative PCR and the 2(-Delta Delta C(T)) Method. Methods. Dec 2001;25(4):402–8. doi:10.1006/meth.2001.1262

40. Kwak S. Are Only. J Lipid Atheroscler. May 2023;12(2):89–95. doi:10.12997/jla.2023.12.2.89

41. Napso T, Lean SC, Lu M, et al. Diet-induced maternal obesity impacts feto-placental growth and induces sex-specific alterations in placental morphology, mitochondrial bioenergetics, dynamics, lipid metabolism and oxidative stress in mice. Acta Physiol (Oxf). Apr 2022;234(4):e13795. doi:10.1111/apha.13795

42. Jones HN, Woollett LA, Barbour N, Prasad PD, Powell TL, Jansson T. High-fat diet before and during pregnancy causes marked up-regulation of placental nutrient transport and fetal overgrowth in C57/BL6 mice. FASEB J. Jan 2009;23(1):271–8. doi:10.1096/fj.08-116889

43. Lopez-Tello J, Sferruzzi-Perri AN. Characterization of placental endocrine function and fetal brain development in a mouse model of small for gestational age. Front Endocrinol (Lausanne). 2023;14:1116770. doi:10.3389/fendo.2023.1116770

44. Caviedes L, Iñiguez G, Hidalgo P, et al. Relationship between folate transporters expression in human placentas at term and birth weights. Placenta. 2016;38:24–28. doi:10.1016/j.placenta.2015.12.007

45. de Barros Mucci D, Kusinski LC, Wilsmore P, et al. Impact of maternal obesity on placental transcriptome and morphology associated with fetal growth restriction in mice. Int J Obes (Lond). 05 2020;44(5):1087–1096. doi:10.1038/s41366-020-0561-3

46. Carter MF, Powell TL, Li C, et al. Fetal serum folate concentrations and placental folate transport in obese women. Am J Obstet Gynecol. Jul 2011;205(1):83.e17–25. doi:10.1016/j.ajog.2011.02.053

47. Chen Y, Zhang J, Cui W, Silverstein RL. CD36, a signaling receptor and fatty acid transporter that regulates immune cell metabolism and fate. J Exp Med. Jun 06 2022;219(6)doi:10.1084/jem.20211314

48. Howell KR, Powell TL. Effects of maternal obesity on placental function and fetal development. Reproduction. 03 2017;153(3):R97–R108. doi:10.1530/REP-16-0495

49. Muscogiuri G, Pugliese G, Laudisio D, et al. The impact of obesity on immune response to infection: Plausible mechanisms and outcomes. Obes Rev. Jun 2021;22(6):e13216. doi:10.1111/obr.13216

50. Inkster AM, Fernández-Boyano I, Robinson WP. Sex Differences Are Here to Stay: Relevance to Prenatal Care. J Clin Med. Jul 05 2021;10(13)doi:10.3390/jcm10133000

51. DiPietro JA, Voegtline KM. The gestational foundation of sex differences in development and vulnerability. Neuroscience. Feb 07 2017;342:4–20. doi:10.1016/j.neuroscience.2015.07.068

52. Broere-Brown ZA, Baan E, Schalekamp-Timmermans S, Verburg BO, Jaddoe VW, Steegers EA. Sex-specific differences in fetal and infant growth patterns: a prospective population-based cohort study. Biol Sex Differ. 2016;7:65. doi:10.1186/s13293-016-0119-1

53. Corrêa RO, Castro PR, Fachi JL, et al. Inulin diet uncovers complex diet-microbiota-immune cell interactions remodeling the gut epithelium. Microbiome. Apr 26 2023;11(1):90. doi:10.1186/s40168-023-01520-2

54. Pellizzon MA, Ricci MR. Choice of Laboratory Rodent Diet May Confound Data Interpretation and Reproducibility. Curr Dev Nutr. Apr 2020;4(4):nzaa031. doi:10.1093/cdn/nzaa031

55. McKeown NM, Fahey GC, Slavin J, van der Kamp JW. Fibre intake for optimal health: how can healthcare professionals support people to reach dietary recommendations? BMJ. Jul 20 2022;378:e054370. doi:10.1136/bmj-2020-054370

56. Seidu Adams CTS, Gui-Xin Qin, Dongsheng Che and Rui Han. Does Dietary Fiber Affect the Levels of Nutritional Components after Feed Formulation? Fibers2018.

57. Meyer U. Prenatal poly(i:C) exposure and other developmental immune activation models in rodent systems. Biol Psychiatry. Feb 15 2014;75(4):307–15. doi:10.1016/j.biopsych.2013.07.011

58. Plociennikowska A, Frankish J, Moraes T, et al. TLR3 activation by Zika virus stimulates inflammatory cytokine production which dampens the antiviral response induced by RIG-I-like receptors. J Virol. Apr 26 2021;95(10)doi:10.1128/JVI.01050-20

